# Training Doctoral Students in Critical Thinking and Experimental Design using Problem-based Learning

**DOI:** 10.1101/2023.03.03.530978

**Authors:** Michael D. Schaller, Marieta Gencheva, Michael R. Gunther, Scott A. Weed

**Affiliations:** Department of Biochemistry and Molecular Medicine, West Virginia University School of Medicine, Morgantown, West Virginia 26506

**Keywords:** Graduate, Training, Critical Thinking, Experimental Design, Problem-Based Learning

## Abstract

**Background:** Traditionally, doctoral student education in the biomedical sciences relies on didactic coursework to build a foundation of scientific knowledge and an apprenticeship model of training in the laboratory of an established investigator. Recent recommendations for revision of graduate training include the utilization of graduate student competencies to assess progress and the introduction of novel curricula focused on development of skills, rather than accumulation of facts. Evidence demonstrates that active learning approaches are effective. Several facets of active learning are components of problem-based learning (PBL), which is a teaching modality where student learning is self-directed toward solving problems in a relevant context. These concepts were combined and incorporated in creating a new introductory graduate course designed to develop scientific skills (student competencies) in matriculating doctoral students using a PBL format.

**Methods:** Evaluation of course effectiveness was measured using the principals of the Kirkpatrick Four Level Model of Evaluation. At the end of each course offering, students completed evaluation surveys on the course and instructors to assess their perceptions of training effectiveness. Pre- and post-tests assessing students’ proficiency in experimental design were used to measure student learning.

**Results:** The analysis of the outcomes of the course suggests the training is effective in improving critical thinking in experimental design. The course was well received by the students as measured by student evaluations (Kirkpatrick Model Level 1). Improved scores on post-tests indicate that the students learned from the experience (Kirkpatrick Model Level 2). A template is provided for the implementation of similar courses at other institutions.

**Conclusions:** A problem-based learning platform appears effective in training newly matriculated graduate students in the required skills for designing experiments to test specific hypotheses, enhancing student preparation prior to initiation of their dissertation research.

## Introduction

For over a decade there have been calls to reform biomedical graduate education. There are two main problems that led to these recommendations and therefore two different prescriptions to solve these problems. The first major issue is the pursuit of non-traditional (nonacademic) careers by doctorates and concerns of adequate training (1, 2). The underlying factors affecting career outcomes are the number of PhDs produced relative to the number of available academic positions (1, 3–5), and the changing career interests of doctoral students (6–9). One aspect in the proposed reformation to address this problem is incorporation of broader professional skills training and creating awareness of a greater diversity of careers into the graduate curriculum (1, 4, 5). Interestingly, a large survey of doctorates in various types of jobs revealed many “soft” or “transferable” skills were equally valued in traditional academic careers as well as non-traditional professions (2). The second issue relates to the curricula content and whether content knowledge, which is continuously expanding, or critical scientific skills should be the core of the curriculum (10, 11). The proposed reformation to address this issue is creation of curricula focusing upon scientific skills, e.g. reasoning, experimental design and communication, while simultaneously reducing components of the curricula that build a foundational knowledge base (12, 13). Components of these two approaches are not mutually exclusive, where incorporation of select specialized expertise in each area has the potential to concurrently address both issues.

One approach that prioritizes the aggregate scientific skill set required for adept biomedical doctorates is the development of core competencies for doctoral students (5), akin to set milestones that must be met by medical residents and fellows (14). Such competencies have been formally established and adopted by multiple top-tier research institutions (15, 16). Key features of these doctoral student competencies include general and field-specific scientific knowledge, critical thinking, experimental design, evaluation of outcomes, scientific rigor, ability to work in teams, responsible conduct of research, and effective communication (5, 15, 16). Such competencies provide clear benchmarks for doctoral committees, graduate programs and higher administration to evaluate the progress of doctoral students’ development into an independent scientific professional and preparedness for the next career stage. Historically, graduate programs relied on traditional content-based courses and supervised apprenticeship in the mentor’s laboratory to develop such competencies. An alternative to this approach is to modify the graduate student curriculum to provide a foundation for these competencies in a more structured way (12, 13). Further, this approach decreases differences that individual graduate students may experience in initial training in these competencies.

One consideration regarding curriculum revision is the learning platform applied to new and revised courses. The relatively new graduate program at the Van Andel Institute Graduate School utilizes a problem-based learning (PBL) platform for delivery of the formal curriculum (17). First developed for medical students (18), the PBL learning approach is now widely used as one component of instruction in many medical schools and has been adopted in other educational settings, including K-12 and undergraduate education (19, 20). A basic tenet of PBL is that student learning is self-directed (18). Students are tasked to solve an assigned problem and are required to find the information necessary for the solution (self-directed). In practice, learning occurs in small groups where a faculty facilitator helps guide the students in identifying gaps in knowledge that require additional study (21). As such, an ideal PBL course is “well organized” but “poorly structured”. The lack of a traditional restrictive structure allows students to pursue and evaluate different solutions to the problem. The premise for PBL is that actively engaging in problem solving enhances learning in several ways (21, 22). First, activation of prior knowledge, as occurs in group discussions, aids in learning by providing a framework to incorporate new knowledge. Second, deep processing of material while learning, e.g. by answering questions or using the knowledge, enhances the ability to later recall key concepts. Third, learning in context, e.g. learning the scientific basis for clinical problems in the context of clinical cases, enables and improves recall. PBL opponents argue that acquisition of large amounts of knowledge, e.g. in the classic medical school classroom setting, is more effective in a traditional didactic curriculum. To directly address this concern, several institutions that changed their medical curricula to a PBL-based format also offered their traditional didactic curriculum in parallel during the transition. Comparison of student performance between the different learning modalities was informative, where students in the PBL-based curriculum performed comparably to the students in traditional curricula on knowledge-centered standardized tests (Step 1 and Step 2). However, PBL-trained students outperformed their peers in the traditional curriculum in clerkships and in interpersonal skills (23, 24), indicating that the PBL-based approach builds a wider platform of practical tools students utilize later in their training. A broader review of medical student performance found that while PBL students did not perform as well on knowledge-based exams, they did slightly better in clinical competencies and outperformed their traditionally-trained peers in interpersonal skills and psychosocial knowledge (25). A comprehensive review of PBL outcomes from K-12 through medical school indicates that PBL students perform better in the application of knowledge and reasoning, but not in other areas like basic knowledge (26). While PBL-trained students recalled less in the short term, long-term knowledge retention was higher than traditional students (21, 26, 27). Although few studies report the outcomes of PBL based approaches in graduate education, this platform may prove useful for training biomedical science doctoral students in critical thinking and practical problem-solving skills.

At our institution, biomedical graduate students enter an umbrella program and take a core curriculum in the first semester prior to matriculating into one of seven biomedical sciences graduate programs across a wide range of scientific disciplines in the second semester. Such program diversity created difficulty in achieving consensus on the necessary scientific foundational knowledge for a core curriculum. Common ground was achieved during a recent curriculum revision through the development of required core competencies for all students regardless of field of study. These competencies and milestones for biomedical science students at other institutions (5, 15, 16) along with nontraditional approaches to graduate education (12, 17), were used as guidelines for curriculum modification. Here we describe the development, implementation and evaluation of a new PBL-based graduate course that provides an initial experience in introducing the scientific career-relevant core competencies of critical thinking and experimental design to incoming biomedical doctoral students.

## Methods

The West Virginia University Institutional Review Board approved the study (WVU IRB Protocol#: 2008081739). Informed consent was provided in writing and all information was collected anonymously. Evaluation of training effectiveness was measured in two ways. First, students completed a questionnaire to capture their perceptions of training upon completion of the course. Second, students took a pre- and post-test to measure differences in their ability to design experiments before and after training. This approach corresponds to the first two levels of the Kirkpatrick Model of Evaluation (28). The pre- and post-tests were identical, asking the students to design an experiment to test a specific hypothesis, include controls, plan analyses, and state possible results and interpretation. Five questions were provided for the pre- and posttest, where each question posed a hypothesis from a different biomedical discipline, e.g. cancer biology or neuroscience. Students were asked to choose one of the five questions to answer. Since answers were anonymous, the analyses compared responses between individuals. Finally, peer-to-peer evaluations were collected to provide feedback on professionalism and teamwork.

All pre- and post-test answers were evaluated by three graders in a blind fashion, where the graders were unaware if an answer came from a pre- or post-test. Prior to grading, each grader made up individual answer keys based upon the question(s) on the tests. The graders then met to compare and deliberate these preliminary keys, incorporating changes and edits to produce a single combined key to use for rating answers. While the students were asked to answer one question, some students chose to answer several questions. Superfluous answers were used as a training dataset for the graders to rate. The graders independently scored each answer, then met to review the results and discuss modification of the grading key. The established final grading key was subsequently utilized by the graders in independently evaluating the complete experimental dataset consisting of all pre- and post-test answers (Appendix VIII).

### Statistical analysis

To measure the interrater reliability of the graders, the intraclass correlation coefficient (ICC) was calculated. A two-way mixed effects model was utilized to evaluate consistency between multiple raters/measurements. The ICC for grading the training dataset was 0.82, which indicates a good inter-rater agreement. The ICC for grading the experimental dataset was also 0.82. For comparison of pre-test vs post-test performance, the scores of the three raters were averaged for each answer. Scores were compared using an unpaired, one-tailed t-test since the average scores for all of the answers on a specific pre- or post-test exhibited a Gaussian distribution and equal variances among the samples. The pre-test and post-test scores for 2020 and 2021 could not be compared due to the different format used for the course in each year.

## Results

The course was created to develop competencies required by all biomedical sciences graduate students regardless of their program of interest (15). As an introductory graduate level course, this met the needs of all our seven diverse biomedical sciences graduate programs where our first-year graduate students matriculate. A PBL platform was chosen for the course to engage the students in an active learning environment (17). The process of problem solving in small teams provided the students with experience in establishing working relationships and how to operate in teams. The students gained experience in researching material from the literature to establish scientific background, find current and appropriate experimental approaches and examples of how results are analyzed. This small group approach allowed each team to develop different hypotheses, experimental plans and analyses based upon the overall interest of the group. The course was designed following discussions with faculty experienced in medical and pharmacy school PBL, and considering course design principles from the literature (19, 29). The specific course objectives are similar to the overall objectives in a graduate program using PBL as the primary course format (Table 1) (17).

**Table 1.**
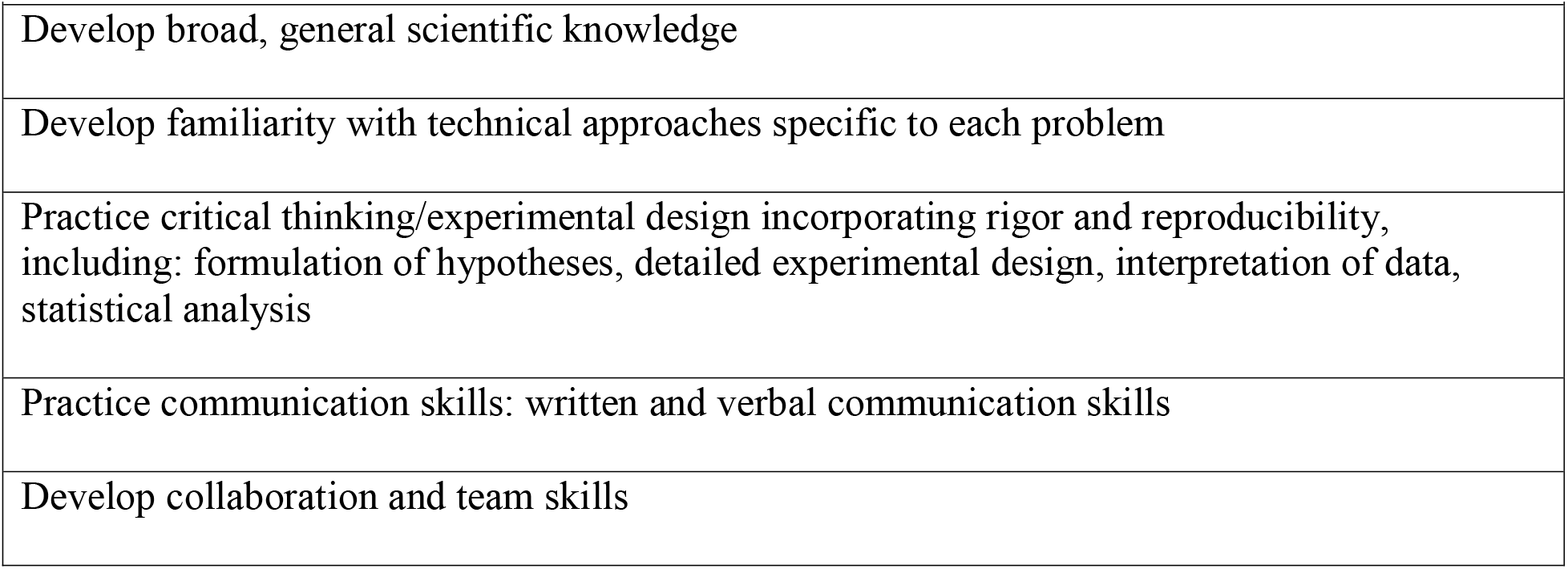
Specific Objectives of the Course.

Students were organized into groups of four or five based on their scientific background. Student expertise in each group was deliberately mixed to provide different viewpoints during discussion. A single faculty facilitator was assigned to each student group, which met formally in 13 separate sessions (Appendix II). In preparation for each session, the students independently researched topics and met informally without facilitator oversight to coordinate their findings and organize the discussion for each class session. During the formal one-hour session, one student served as the group leader to manage the discussion. The faculty facilitator guided the discussion to ensure coverage of necessary topics and helped the students identify learning issues, i.e. areas that require additional development, for the students to research and address for the subsequent session. At the end of each session, teams previewed the leading questions for the following class and organized their approach to address these questions prior to the next session.

As the course was developed during the COVID-19 pandemic, topics related to SARS-CoV2 and COVID-19 were selected as currently relevant problems in society. Session 1 prepared the students to work in teams by discussing about how to work in teams and manage conflict. In session 2, the students met in their assigned groups to get to know each other, discuss problem-based learning and establish ground rules for the group. Sessions 3 and 4 laid the course background by focusing on the SARS-CoV2 virus and COVID-19-associated pathologies. The subsequent nine sessions were organized into three separate but interrelated three-session blocks: one on COVID-19 and blood clotting, one on COVID-19 and loss of taste, and one on SARS-CoV2 and therapeutics. The first session in each of these blocks was devoted to covering background information (blood clotting, neurosensation and drug application). The second session of each block discussed hypothesis development (mechanisms that SARS-CoV2 infection might utilize to alter blood clotting, the sense of taste, and identification of therapeutic targets to attenuate SARS-CoV2 infection). In the second sessions the students also began to design experiments to test the hypothesis. The final session of each block fleshed out the details of the experimental design.

The process was iterative, where the students had three opportunities to discuss hypothesis development, experimental design and analysis during sessions with their facilitators. Written and oral presentation assignments (Appendix V) provided additional opportunities to articulate a hypothesis, describe experimental approaches to test the hypotheses and propose analysis of experimental results.

Rigor and reproducibility was incorporated into the course. As the students built the experimental design to address the hypothesis, recurring questions were posed to encourage them to consider rigor. Examples include: “*Are the methods and experimental approaches rigorous? How could they be made more rigorous?”* “*Discuss how your controls validate the outcome of the experiment. What additional controls could increase confidence in your result?”* The facilitators were instructed to direct discussion to topics related to the rigor of the experimental design. The students were asked about numbers of replicates, number of animals, additional methods that could be applied to support the experiment, and other measurements to address the hypothesis in a complementary fashion. In the second iteration of the course, we introduced an exercise on rigor and reproducibility for the students using the NIH Rigor and Reproducibility Training Modules (see Appendix III). In this exercise, the students read a short introduction into rigor and reproducibility and viewed a number of short video modules to introduce lessons on rigor. The students were also provided the link to the NIGMS clearinghouse of training modules on rigor and reproducibility as reference for experimental design in their future (see Appendix III).

The first delivery of the course was during the COVID-19 pandemic and sessions were conducted on the Zoom platform. The thirteen PBL sessions, and two additional sessions dedicated to oral presentations, were spaced over the course of the first semester of the biomedical sciences graduate curriculum. The second iteration of the course followed the restructuring of the graduate first year curriculum and the thirteen PBL sessions, plus one additional session devoted to oral presentations, were held during the first three and a half weeks of the first-year curriculum. During this period in the semester, this was the only course commitment for the graduate students. Due to this compressed format, only one written assignment and a single oral presentation were assigned. As the small group format worked well via Zoom in the first iteration of the course, the small groups continued to meet using this virtual platform.

At the beginning of each year, a pre-test was administered to assess the ability of the incoming students to design experiments. Students were asked to answer the following question: *“Design an experiment to test the hypothesis that____________. Describe your experiment(s), necessary controls and planned analysis. Describe the possible results of your experiment(s) and how these results will be interpreted.”* Students were given five separate questions from different biomedical fields that were structured in an identical manner. Each question had a unique hypothesis to test: 1. protein X controls body weight, 2. overexpression of protein X causes cancer, 3. a kinase, protein X, phosphorylates protein Y, 4. an anti-oxidant (Vitamin C) effects vascular health with aging, and 5. protein X controls adult neurogenesis. Students were tasked with answering any one of the five questions and thus had the choice of topic. Twenty-six students completed the pre-test in each year. To determine if an improvement in experimental design could be measured after taking the course, the identical test was given following course completion as a post-test. Eighteen students completed the post-test at the end of the first year and 26 students completed the test at the end of the second year. The higher response rate in the second year might be due to fewer end of semester surveys since this was the only course that the students took in that time period. Additionally, the post-test in the second year was conducted at a scheduled time, rather than on the student’s own time as was the case in year one. Pre- and post-tests were conducted anonymously each year in an effort to maximize survey participation. Question selection (excluding students that misunderstood the assignment and answered all questions) is shown in Table 2. The most frequently selected questions were Question 1 (45 times) and Question 2 (23 times). Interestingly, the results in Table 2 also indicate that students did not necessarily choose the same question to answer on the pre-test and post-test.

**Table 2.**
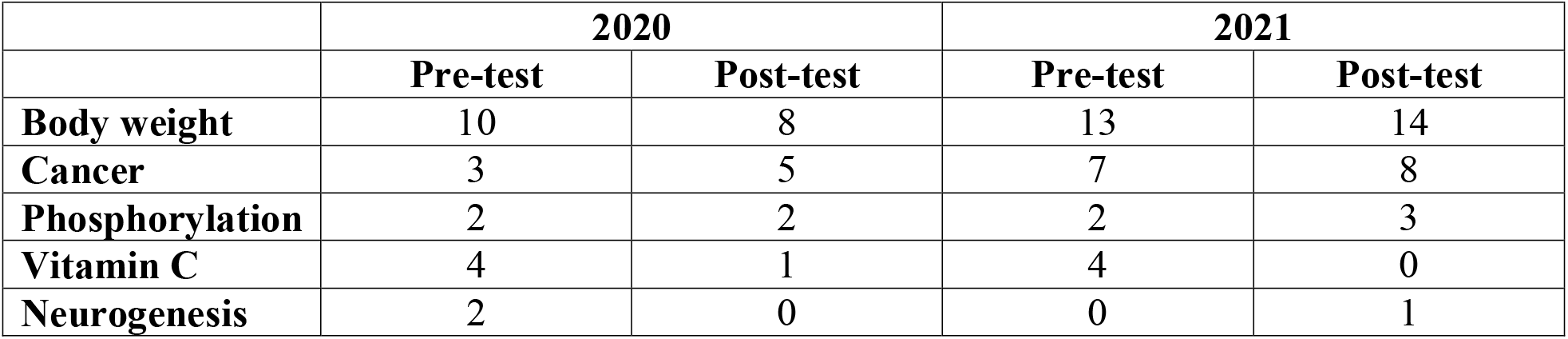
Student Choice of Experimental Question to Answer (Only those who made a choice)

|  |  |  |  |  |
| --- | --- | --- | --- | --- |
|  | **2020** |  | **2021** |  |
|  | **Pre-test** | **Post-test** | **Pre-test** | **Post-test** |
| **Body weight** | 10 | 8 | 13 | 14 |
| **Cancer** | 3 | 5 | 7 | 8 |
| **Phosphorylation** | 2 | 2 | 2 | 3 |
| **Vitamin C** | 4 | 1 | 4 | 0 |
| **Neurogenesis** | 2 | 0 | 0 | 1 |

At the end of the course, students were asked to respond to surveys evaluating the course and their facilitator (see Appendix VII). In the first year, 23 students responded to the course evaluation and 26 students submitted facilitator evaluations, whereas in the second year there were 25 responses to the course evaluation and 26 for facilitators. A peer-to-peer survey was also utilized with a Goldilocks scale to assess professionalism and colleague interactions (see Appendix VII). In the second course iteration, Goldilocks surveys were collected three times over the three-week course period due to the compressed time frame. This was necessary to provide rapid feedback to the students about their performance during the course in order provide opportunities to address and rectify any deficits before making final performance assessments.

### Student reception of the course

Likert scores for the 2020 and 2021 course evaluations are presented in Figure 1. The median score for each question was 4 on a scale of 5 in 2020. In 2021, the median scores for the questions about active learning and hypothesis testing were 5 and the median score of the other questions was 4. The students appreciated the efforts of the facilitators in the course, based upon their evaluations of the facilitators. The median score for every facilitator across all survey questions is shown in Figure 2. The median score for a single question in 2020 and 2021 was 4.5 and the median score for all other questions was 5. The results of the peer-to-peer evaluations are illustrated in Figure 3. A Goldilocks scale ranging from 1 to 7 was used, with 4 being the desired score. An example peer question asked about accountability, where responses included *not accountable, e.g. always late* (score = 1), *accountable, e.g punctual, well prepared, follows up* (score = 4) and *controlling, e.g. finds fault in others* (score =7). Each student provided a peer-to-peer evaluation for each student in their group. The average score for each student were plotted, with scores further from the desired score of 4 indicating perceived behaviors that were not ideal. The wide range of scores in the 2020 survey were noted. The students completed three peer-to-peer surveys during the 2021 course. The range of scores in the 2021 peer-to-peer evaluation was narrower than the range in the 2020 survey. The range of scores was expected to narrow from the first (initial) to third (final) survey as students learned and implemented improvements in their professional conduct based upon peer feedback. The narrow range of scores in the initial survey left little room for improvement.

**Figure 1.**
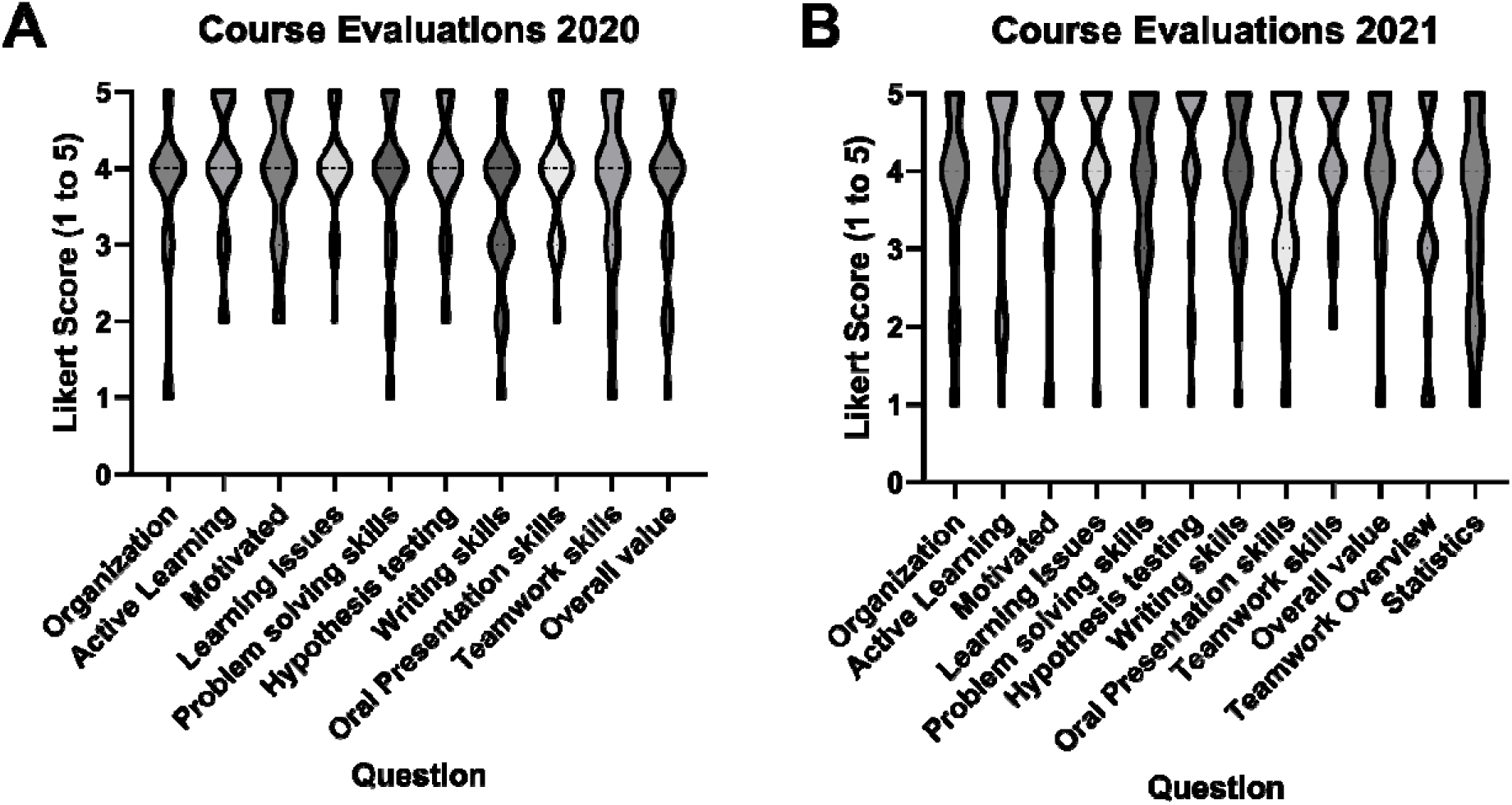
Results of Course Evaluations by Students. Student evaluations of the course were collected at the end of each offering. The evaluation surveys are in Appendix VII. Violin plots showing the distribution and median score for each question in the 2020 survey **(A)** and the 2021 survey **(B)** are shown. The survey used a Likert scale (1 – low to 5 – high).

**Figure 2.**
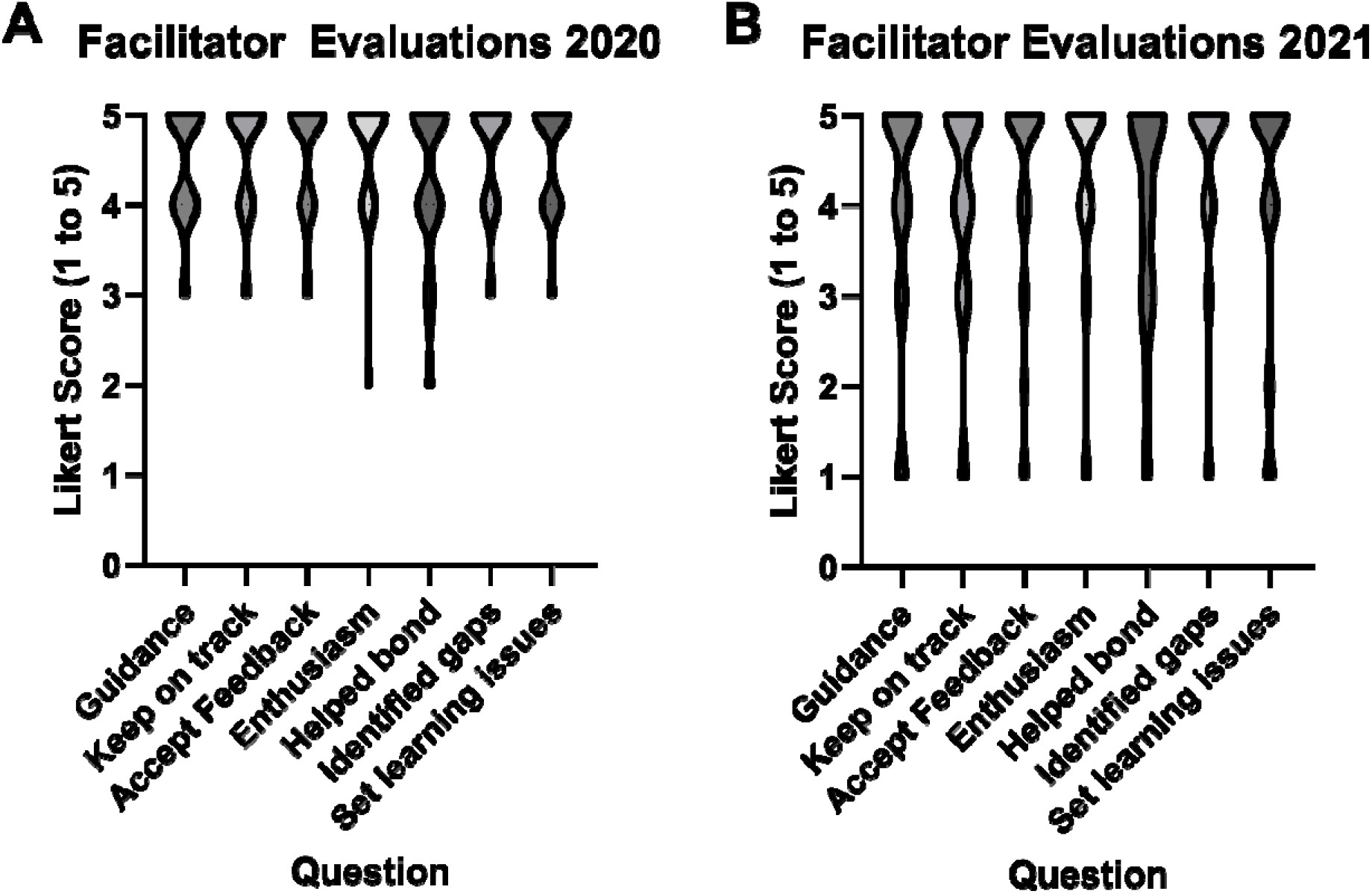
Results of Facilitator Evaluations by Students. Student evaluations of the facilitators were collected at the end of each offering of the course. The evaluation surveys are in Appendix VII. Violin plots showing the distribution and median score for each question in the 2020 survey **(A)** and the 2021 survey **(B)** are shown. The survey used a Likert scale (1 – low to 5 – high).

**Figure 3.**
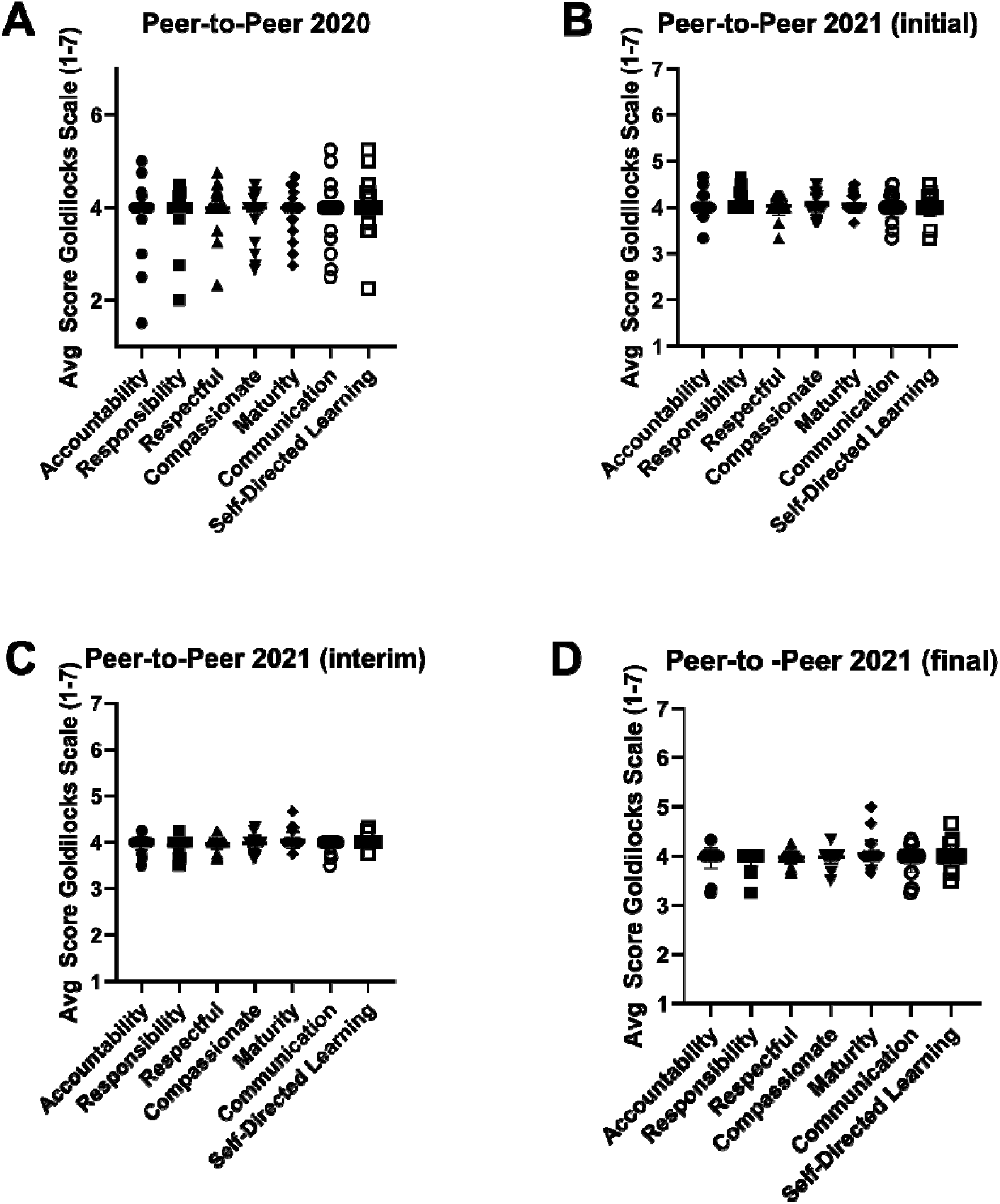
Results of Student Peer-to-Peer Evaluations. Student peer-to-peer evaluations were collected at the end of the course in year 1 **(A)**, and at the beginning **(B)**, the middle **(C)** and the end **(D)** of the course in year 2. Each student evaluated the professionalism of each other student in their group using the evaluation survey shown in Appendix VII. The average score for each student is plotted as a data point. The survey used a Goldilocks scale (range of 1 to 7) where the desired professional behavior is reflected by a score of 4.

### Student learning

To assess the ability of student cohorts to design experiments before and after the course, three of the authors graded all of the pre- and post-test answers. Grading was performed in a blind fashion, where answers could not be identified as a pre-test or post-test answer. A common grading key was used by the three raters, where 16 was a perfect answer score (Appendix VIII). The scores of the three raters were averaged for each answer. The intraclass correlation coefficient of the combined scores across both years was 0.82, indicating a good inter-rater agreement. Average scores on each test were Gaussian distributed and exhibited similar variances, permitting further parametric analysis. Since the format of the course was different in the two iterations, comparison of test results between the two years was not performed. The average scores of the pre- and post-test in 2020 were not statistically different (p ~ 0.07), although the post-test scores trended higher. In contrast, the difference between the pre- and post-test in 2021 did reach statistical significance (p~0.03). The results collectively indicate an overall improvement in student ability in experimental design (Figure 4).

**Figure 4.**
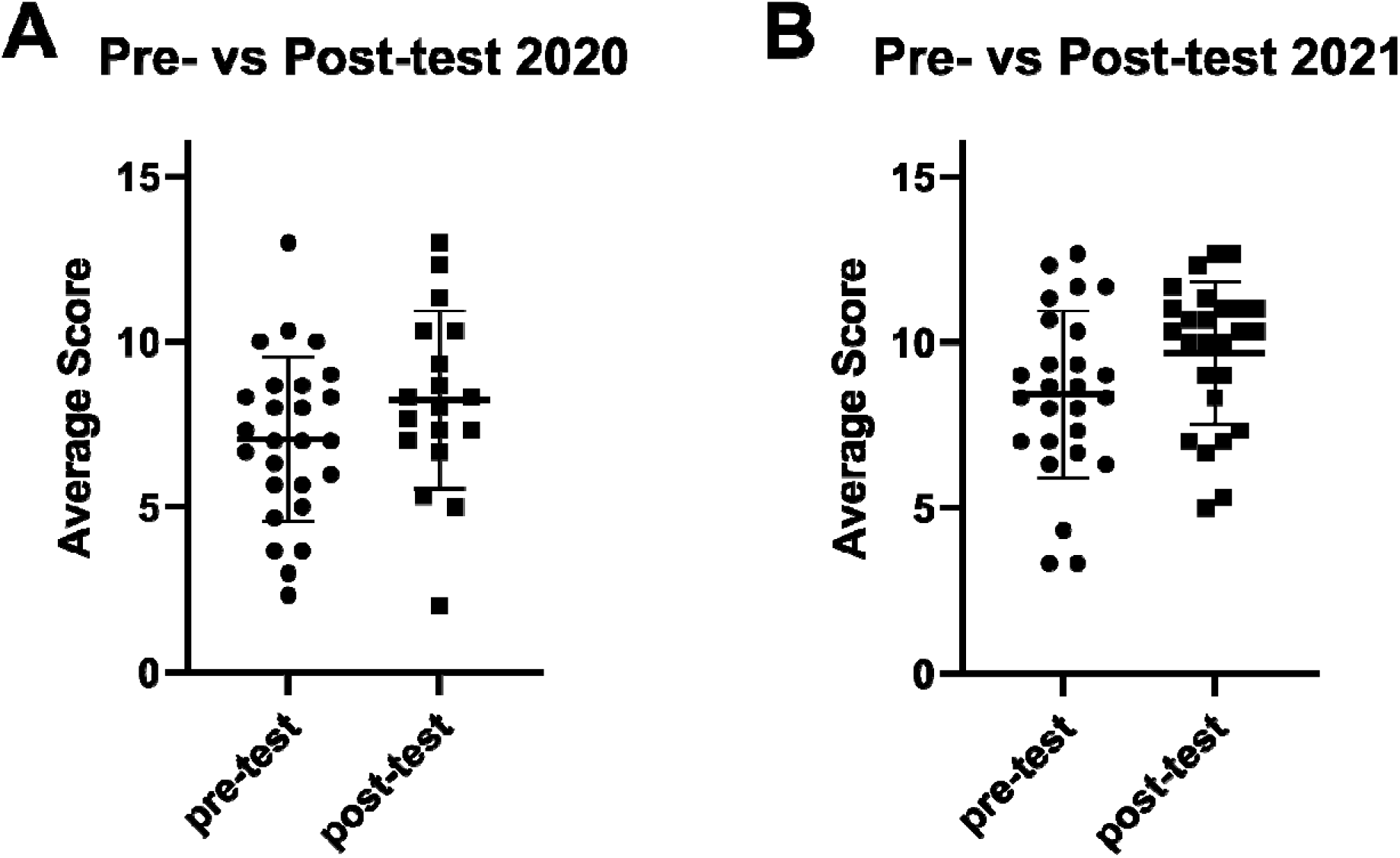
Pre- and Post-Test Scores. At the beginning and end of each offering, the students completed a test to measure their ability to design an experiment (see text for the details of the exam). Three faculty graded every answer to the pre- and post-test using a common grading rubric (see Appendix VIII). The maximum possible score was 16. The average score for each individual answer on the pre-test and post-test is represented as a single data point. The bar indicates the mean score across all answers +/- SD. The average scores of the pre- and post-test scores were compared using an unpaired, one-tailed t-test. For the 2020 data **(A)**, p = 0.0695, and for the 2021 data **(B)**, p = 0.0294.

## Discussion

Here we detail the creation, implementation and assessment of a first-year biomedical sciences graduate course that provides training in the core graduate student competencies of experimental design, communication and teamwork using a PBL platform. Student evaluations indicated the course was effective at motivating active learning and that students became more active learners. The evaluation survey questions were directly related to three specific objectives of the course: 1) Students reported developing skills in problem solving, hypothesis testing and experimental design. 2) The course helped develop oral presentation skills and written communication skills (in one iteration of the course) and 3) students developed collaboration and team skills. Thus, from the students’ perspective, these three course objectives were met. Student perceptions of peer professionalism was measured using peer-to-peer surveys. The wide range of Goldilocks scores in the first student cohort was unexpected. In the second student cohort changes in professional behavior were measured over time and the score ranges were narrower. The reasons for the difference between cohorts is unclear. One possibility is that the first iteration of the course extended over one semester and was during the first full semester of the pandemic, impacting professional behavior and perceptions of professionalism. The second cohort completed a professionalism survey three times during the course. The narrow range of scores from this cohort in the initial survey made detection of improved professionalism over the course difficult. Results do indicate that professionalism improved in terms of respect and compassion between the first and last surveys. Finally, the results of the pre-test and post-test analysis demonstrated a trend of improved performance on the post-test relative to the pre-test for students in each year of the course and a statistical difference between the pre- and post-test scores in the second year.

### Areas for improvement

The course was initially offered as a one-credit course. Student comments on course evaluations and comments in debriefing sessions with facilitators at the end of the course concurred that the work load exceeded that of a one credit course. As a result, the year two version was offered as a two-credit course to better align course credits with workload.

There were student misperceptions about the goals of the course in the first year. Some students equated experimental design with research methods and expressed disappointment that this was not a methods course. While learning appropriate methods is a goal of the course, the main emphasis is developing hypotheses and designing experiments to test the hypotheses. As such, the choice of methods was driven by the hypotheses and experimental design. This misperception was addressed in the second year by clearly elaborating on the course goals in an introductory class session.

The original course offering contained limited statistical exercises to simulate experimental planning and data analysis, e.g. students were required to conduct a power analysis. Between the first and second years of the course, the entire first semester biomedical sciences curriculum was overhauled with several new course offerings. This new curriculum contained an independent biostatistics workshop that students completed prior to the beginning of this course. Additional statistics exercises were incorporated into the PBL course to provide the students with more experience in the analysis of experimental results. Student evaluations indicated that the introduction of these additional exercises was not effective. Improved coordination between the biostatistics workshop and the PBL course is required to align expectations, better equipping students for the statistical analysis of experimental results encountered later in the course.

An important aspect that was evident from student surveys, facilitator discussions and debrief sessions was that improved coordination between the individual facilitators of the different groups is required to reduce intergroup variability. Due to class size, the students were divided into six groups, with facilitators assigned to the same group for the duration of the course to maintain continuity. The facilitators met independent of the students throughout the course to discuss upcoming sessions and to share their experiences with their respective groups. This allowed the facilitators to compare approaches and discuss emerging or perceived concerns/issues. In the second year, one facilitator rotated between different groups during each session to observe how the different student groups functioned. Such a real time faculty peer-evaluation process has the potential to reduce variability between groups, but was challenging to implement within the short three-week time period. Comprehensive training where all facilitators become well versed in PBL strategies and adhere to an established set of guidelines/script for each session is one mechanism that may reduce variability across different facilitator-group pairings.

### Limitations

The current study has a number of limitations. The sample size for each class is small. Due to restructuring of the graduate curriculum and the pandemic, the two iterations of the course were formatted differently. This precluded pooling the data from the two offerings and makes comparison between the outcomes difficult. Six different PBL groups were required to accommodate the number of matriculating students in each year. Despite efforts to provide a consistent experience, there was variability between the different facilitators in running their respective groups. This could be responsible for increasing the spread of scores on the post-tests and decreasing the value of the course for a subset of students. The pre- and post-tests were conducted anonymously to encourage student participation. This prevented correlating the differential between pre- and post-test scores for each student and in comparing learning between different groups. While the course analysis captured the first two levels of the Kirkpatrick model of evaluation (reaction and learning), it did not attempt to measure the third level (behavior) or fourth level (results) (28). Future studies are required to measure the third level. This could be achieved by asking students to elaborate on their experimental design used in recent experiments in their dissertation laboratory following completion of the course, or by evaluating the experimental design students incorporate into their dissertation proposals. The fourth Kirkpatrick level could potentially be assessed by surveying preceptors about their students’ abilities in experimental design in a longitudinal manner at semi- or annual committee meetings and accompanying written progress reports. The advantage of focusing on the first two Kirkpatrick levels of evaluation is that the measured outcomes can be confidently attributed to the course. Third and fourth level evaluations are more complicated, since they necessarily take place at some point after completion of the course. Thus, the third and fourth level outcomes can result from additional factors outside of the course (e.g. other coursework, working in the lab, attendance in student-based research forum, meeting with mentors, etc.). Another limiting factor is the use of a single test to measure student learning. Additional alternative approaches to measure learning might better capture differences between the pre- and post-test scores.

### Implementation

Development of an effective PBL course takes considerable time and effort to conceive and construct. Successful implementation requires the requisite higher administrative support to identify and devote the necessary and appropriate faculty needed for course creation, the assignment of skilled faculty to serve as facilitators and staff support to coordinate the logistics for the course. It is critical that there is strong faculty commitment amongst the facilitators to devote the time and energy necessary to prepare and to successfully facilitate a group of students. Strong institutional support is linked to facilitator satisfaction and commitment to the PBL-based programs (30). Regular meetings between the course coordinator and facilitators to discuss the content of upcoming sessions and define rubrics to guide student feedback and evaluation were mechanisms used to standardize between the different groups in this course (Appendix VI). In hindsight, the course would benefit from more rigorous facilitator training prior to participation in the course. While a number of our facilitators were veterans of medical school PBL course, the necessary skillset required to effectively manage a graduate level PBL course that is centered on developing critical thinking and experimental design are different. Such training requires an extensive time commitment by the course coordinators and participating facilitators, a factor that should be recognized during annual faculty evaluations by departmental chairs and promotion and tenure committees.

The most difficult task in developing this course involved the course conception and development of the problem-based assignments. Designing a COVID-19 based PBL course in 2020 required *de novo* development of all course material. This entailed collecting and compiling information about the virus and the disease to provide quick reference for facilitators to guide discussion in their groups, all in the face of constantly shifting scientific and medical knowledge, along with the complete lack of traditional peer-based academic social engagement due to the pandemic. In development of this course, three different COVID-based problems were identified, with appropriate general background material for each problem requiring extensive research and development. Background material on cell and animal models, general strategies for experimental manipulation and methods to measure specific outcomes were collected in each case. Student copies for each session were designed to contain a series of questions as a guide to identifying important background concepts. Facilitator copies for each session were prepared with the goal of efficiently and effectively guiding each class meeting. These guidelines contained ideas for discussion points, areas of elaboration and a truncated key of necessary information to guide the group (Appendix IV). Several PBL repositories exist (e.g. https://itue.udel.edu/pbl/problems/, https://www.nsta.org/case-studies) and MedEdPORTAL (https://www.mededportal.org/) publishes medical-specific cases. These provide valuable resources for case-based ideas, but few are specifically geared for research-focused biomedical graduate students. As such, modification of cases germane to first year biomedical graduate students with a research-centered focus is required prior to implementation. Finally, appropriate support materials for surveys and evaluation rubrics requires additional development and refinement of current or existing templates to permit improved evaluation of learning outcomes (Appendix VI).

### Future Directions

Novel curriculum development is an often overlooked but important component to contemporary graduate student education in the biomedical sciences. It is critical that modifications incorporated in graduate education are evidence based. We report the implementation of a novel PBL course for training in the scientific skill sets required for developing and testing hypotheses, and demonstrate its effectiveness. Additional studies are required to determine the long-term impact of this training on student performance in the laboratory and progression towards degree. It will be interesting to determine if similar curriculum changes to emphasize development of skills will shorten the time to degree, a frequent recommendation for training the modern biomedical workforce (1, 31–33). Incorporation of courses emphasizing development of skills can be done in conjunction with traditional didactic instruction to build the necessary knowledge base for modern biomedical research. Our PBL course was stand-alone, requiring the students to research background material prior to hypothesis development and experimental design. Coordination between the two modalities would obviate the need for background research in the PBL component, reinforce the basic knowledge presented didactically through application, and prepare students for higher order thinking about the application of the concepts learned in the traditional classroom. Maintaining a balance between problem-based and traditional instruction may also be key in improving faculty engagement into such new and future initiatives. Continued investments in the creation and improvement of innovative components of graduate curricula centered around developing scientific skills of doctoral students can be intellectually stimulating for faculty and provide a better training environment for students. The effort may be rewarded by streamlining training and strengthening the biomedical workforce of the future.

## Abbreviations

PBL: problem-based learning
ICC: intraclass coefficient
SARS-CoV2: severe acute respiratory syndrome coronavirus 2
COVID-19: coronavirus disease 19

## Acknowledgements

Thanks to Mary Wimmer and Drew Shiemke for many discussions over the years about PBL in the medical curriculum and examples of case studies. We thank Steve Treisenberg for initial suggestions and discussions regarding PBL effectiveness in the Van Andel Institute. Thanks to Paul and Julie Lockman for discussions about PBL in School of Pharmacy curricula and examples of case studies. Special thanks to the facilitators of the groups, Stan Hileman, Hunter Zhang, Paul Chantler, Yehenew Agazie, Saravan Kolandaivelu, Hangang Yu, Tim Eubank, William Walker, and Amanda Gatesman-Ammer. Without their considerable efforts the course could never have been successfully implemented. Thanks to the Department of Biochemistry and Molecular Medicine for supporting the development of this project. MS is the director of the Cell & Molecular Biology and Biomedical Engineering Training Program (T32 GM133369).

## Authors’ contributions

SW and MS developed the concept for the course. MS was responsible for creation and development of all of the content, for the implementation of the course, the design of the study and creating the first draft of the manuscript. MG, MRG and SW graded the pre- and post-test answers in a blind fashion. MS, MG, MRG and SW analyzed the data and edited the manuscript.

## Funding

There was no funding available for this work.

## Availability of data and materials

All data generated in this study are included in this published article and its supplementary information files.

## Declarations

### Ethics approval and consent to participate

The West Virginia University Institutional Review Board approved the study (WVU IRB Protocol#: 2008081739). Informed consent was provided in writing and all information was collected anonymously. All methods were carried out in accordance with relevant guidelines and regulations.

### Consent for publication

Not applicable.

### Competing interests

The authors declare no financial or non-financial competing interests.

## Notes

### Competing Interest Statement

The authors have declared no competing interest.

