## Supplemental Material for "Training Doctoral Students in Critical Thinking and Experimental Design using Problem-based Learning"

**Table of Contents**

APPENDIX I – Course Syllabus	pg 2

APPENDIX II – Session Topics	pg 5

APPENDIX III – Rigor and Reproducibility	pg 6

APPENDIX IV – Facilitators’ Copy for each Session	pg 7

Session 2 – Welcome	pg 7

Session 3 – The Virus	pg 9

Session 4 – The Pathology	pg 17

Session 5 – Cardiovascular/Clotting Problems	pg 25

Session 6 – Hypothesis Development	pg 31

Session 7 – Experimental Design	pg 35

Session 8 – Neurological Problems	pg 37

Session 9 – Hypothesis Development	pg 45

Session 10 – Experimental Design	pg 49

Session 11 – SARS-CoV2 Therapeutics	pg 51

Session 12 – Hypothesis Development	pg 55

Session 13 – Experimental Design	pg 57

APPENDIX V – Assignments	pg 59

APPENDIX VI – Rubrics	pg 61

Class Participation	pg 61

Written Report	pg 62

Oral Presentation	pg 64

Peer to Peer Evaluation	pg 67

APPENDIX VII – Evaluation Surveys	pg 69

APPENDIX VIII – Grading Rubric	pg 72

###### APPENDIX I COURSE SYLLABUS

###### Critical Thinking and Experimental Design

---------------------------------------------------------------------------------------------------------------------

**Course Objectives:**

Purpose of the Biomedical Research Methods course is development of competencies that will be useful in all graduate programs and all laboratories. The platform for delivery is a current or historically important medical problem. Students work in teams to develop competencies while addressing the problem.

Specific Objectives

- Develop broad, general scientific knowledge
- Develop familiarity with technical approaches specific to each problem
- Practice critical thinking/experimental design incorporating rigor and reproducibility, including:
  - Formulation of hypotheses
  - Detailed experimental design
  - Interpretation of data
  - Statistical analysis
- Practice communication skills, including:
  - Written communication skills
  - Oral communication skills
- Develop collaboration and team skills

Specific Learning Outcomes

- Demonstrate general scientific knowledge related to the problems discussed
- Utilize appropriate technical approaches to address novel problems in areas related to problems addressed in class
- Design rigorous experiments directly addressing novel problems, analyze and interpret results
- Effectively articulate scientific concepts, approaches and outcomes of experiments in written reports and oral reports
- Demonstrate collaborative team skills

---------------------------------------------------------------------------------------------------------------------

**Expectations of Students:**

- Work together as a team to solve problems
- Establish a positive environment for discussion and encouraging open communication
- Participate fully in discussion
- Respect one another and show kindness by listening, understanding the diverse perspective of others and responding in a constructive way
- Work individually to complete written assignments, drawing upon information compiled by the team
- Work as a team to develop oral presentations

---------------------------------------------------------------------------------------------------------------------

**Evaluation:**

Student evaluations will be based on participation in classroom discussions, teamwork, written and oral reports on problems (see rubrics). The course is graded pass/fail.

---------------------------------------------------------------------------------------------------------------------

**Attendance policy:**

Students are expected to attend all sessions. If a student must miss a class, they must communicate with the instructor and their group PRIOR to the class to receive an excused absence. Absences due to emergency must be communicated with the instructor as soon as possible, prior to class, if possible. Accumulation of unexcused absences will affect the student’s final grade.

---------------------------------------------------------------------------------------------------------------------

**Social Justice Statement:**

The West Virginia University community is committed to creating and fostering a positive learning and working environment based on open communication, mutual respect, and inclusion. Any suggestions as to how to further such a positive and open environment in this class will be appreciated and given serious consideration.

If you are a person with a disability and anticipate needing any type of accommodation in order to participate in this class, please advise the coordinator and make appropriate arrangements with the [Office of Accessibility Services](http://diversity.wvu.edu/oas)(304-293-6700). For more information on West Virginia University’s Diversity, Equity, and Inclusion initiatives, please see http://diversity.sandbox.wvu.edu

**APPENDIX II SESSION TOPICS**

**Critical Thinking and Experimental Design**

**Session Topics**

1 - Working in Teams (meet as a class – all other sessions meet in groups)

2 – Initial group meeting – ice breaking – establishing ground rules

3 - SARS-CoV2 - the virus

4 - SARS-CoV2 - pathogenesis

5 - COVID-19 and coagulation

6 - COVID-19 and coagulation - hypotheses

7 - COVID-19 and coagulation - expt. design

8 - COVID-19 and taste

9 - COVID-19 and taste - hypothesis

10 - COVID-19 and taste - expt. design

11 - SARS-CoV2 therapeutics

12 - SARS-CoV therapeutics - hypothesis

13 - SARS-CoV2 therapeutics - expt. Design

**APPENDIX III RIGOR AND REPRODUCIBILITY**

**Rigor and Reproducibility**

The scientific community is concerned about the rigor and reproducibility of research. The rigor of research must be addressed in grant applications to the NIH and other agencies. Journals are also scrutinizing the rigor of research prior to publication. To better prepare trainees, curricula are being modified to incorporate rigor and reproducibility. The National Institutes of Health has prepared several training modules to illustrate some aspects of rigorous experimental design.

Please read the PDF file Introduction to the NIH Modules. It is only two pages and mostly bullet points. There are a series of questions at the end of the document. Think about these when you are designing experiments in this course.

Please view these four YouTube videos from the NIH to introduce yourself to some concepts relevant to scientific rigor (total viewing time is only ~15 minutes). The videos are a little corny, but they make a point.

Module 1 – Lack of Transparency - <https://www.youtube.com/watch?v=idt7_GWd088>

Module 2 – Blinding and Randomization - <https://www.youtube.com/watch?v=cOVRUjNk6u8>

Module 3 – Biological and Technical Replicates - <https://www.youtube.com/watch?v=gBntka4O22Y>

Module 4 – Sample Size, Outliers, and Exclusion Criteria - <https://www.youtube.com/watch?v=pKVKeO9pB0s>

The National Institute of General Medical Sciences maintains a clearinghouse of training modules on various topics related to rigor and reproducibility. <https://www.nigms.nih.gov/training/pages/clearinghouse-for-training-modules-to-enhance-data-reproducibility.aspx>

The materials available in this clearinghouse may be useful to you as you advance in your graduate career and incorporate different types of experiments into your project.

**APPENDIX IV FACILITORS’ COPY FOR EACH SESSION**

The content of the facilitators’ copy of each session is included. The content of the students’ copy for each session is demarcated in italics and underlined. Note that session 1 was a didactic presentation about working in teams.

**SESSION 2 – Welcome**

*Introduction*

- Ice breaker/getting to know you activity, e.g. creating a group biography – things like hometowns, experiences, career goals, favorite things (courses, hobbies) (1)

Using groups as a learning strategy

- What benefits do you, the facilitator, see in using groups as a learning strategy?

*What experiences do you have in using groups as a learning strategy?*

- Facilitate the discussion making sure all who want to share have the opportunity
- What were their good experiences? Bad experiences? (**Facilitator has to take notes in this session**)

Discuss working in teams

- Perhaps some activity to illustrate benefits of solving problems in teams (1)
  - E.g., one student is ‘teacher’. Show teacher a diagram with multiple shapes (~3) each with a specific orientation. Teacher describes to students, who can take notes (but not draw) – **important 2 minute limit**
  - Students draw the picture – 2 minute limit
  - Students discuss their pictures and come up with a consensus picture – 5 minute limit
  - Compare individual pictures with consensus – teacher judges the most accurate

*What are the attributes of a good team player?*

- Facilitate discussion and take notes

*What are the attributes of a poor team player?*

- Facilitate discussion and take notes

*What should our ground rules be for the group?*

- Facilitate discussion and take notes on establishing ground rules for the group – enforce these as a team, e.g.
- Be on time
- Be prepared
- Notify all if you need to miss a class
- Respect the views, values and ideas of other members
- Also need to have consequences

*We will assign rotating roles for group members. The facilitator will discuss.*

Group roles:

- Discussion leader – keeps on track – maintains full participation
- Recorder – records bullets of discussion, shares screen to recap bullets at the end
- Scribe – records assignments, strategies, learning issues, data
- Researcher – finds definitions for unknown terms and clarifies jargon

Refs

1.	Duch BJ, Groh SE, Allen DE. The power of problem-based learning: a practical" how to" for teaching undergraduate courses in any discipline: Stylus Publishing, LLC.; 2001.

**SESSION 3 -Coronavirus problem – the virus**

*In December 2019, patients were admitted to hospitals in Wuhan with a pneumonia of unknown etiology. This disease, called COVID-19, was caused by a novel coronavirus, called SARS-CoV2. In the twenty months following the original emergence of the virus in the human population,* there have been 205,611,815 cases of infection worldwide resulting in 4,338,593 deaths. Johns Hopkins University reports 36,307,177 infections in the US resulting in 619,094 deaths (as of Aug 13, 2021) (<https://coronavirus.jhu.edu/map.html>).

*Understanding the virus and its life cycle in human cells is critical to understanding the disease and developing therapeutic strategies.*

*Describe the SARS-CoV2 virus.*

***Some Possible Discussion Topics***

- What other viruses are related to SARS-CoV2 and with what diseases are they associated?
- What is the structure and organization of the genome?
- What is the difference between a +ve stranded virus and a -ve stranded virus?
- What proteins are found in the virus?
- What are the functions of these proteins (particularly the spike protein, S)?
- What are the lipid components in the virus?
- What problem does the lipid pose in the infection process?

*How does SARS-CoV2 get into a human cell?*

***Some Possible Discussion Topics***

- How does the virus attach to the cell – what is the receptor on the virus, what is the receptor on the cell?
- What is the normal function of the receptor on the surface of the cell?
- How is the receptor on the virus (S) modified to promote its function of cell attachment – what enzyme is responsible?
- Where does the viral genome need to go in the cell – where does translation take place?
- How does the viral genomic material get delivered to the cytoplasm?
- What viral protein is responsible for promoting viral fusion?
- How is this protein, the S2 fragment of the spike protein, modified to promote its function of membrane fusion?

*What events occur during viral replication in a human cell?*

***Some Possible Discussion Topics***

- How are new viral proteins made?
- What are the differences between non-structural proteins and structural proteins?
- How are viral mRNAs and new viral genomes made?
- What viral proteins are required for transcription and replication?
- How are the virus components assembled into new virus particles?
- How is the virus released from the cell?

*Since the isolation of the original virus, a number of variants have emerged. Using one variant as an example, discuss the general significance of alterations in the virus.*

- Which genes contain pathologically significant mutations in the SARS-CoV2 variants?
- Why would you expect these mutations to alter transmission?
- Why would you expect these mutations to alter the response to antibody therapy or resistance to vaccination?
- How could these mutations alter detection in a SARS-CoV2 test?

**xxxxxxxxxxxxxxxxxxxxxxxxxxxxxxxxxxxxxxxxxxxxxxxxxxxxxxxxxxxxxxxxxxxxxxxxxxxxxxxxxxxx**

***Facilitator Cheat Sheet***

**Coronavirus problem, SARS Cov2 bullets**

**Genetic material – SARS-CoV2**

- RNA
- Single stranded
- +ve stranded, i.e. genome can be translated
- 30,000 base genome
- 5’ cap and polyA tail
- 12-14 coding regions
- 5 major genes – Replicase (Nsp polyprotein – 16 predicted nonstructural proteins), Spike (S), Envelope (E), Membrane (M), Nucleocapsid (N)
- Nsps include RNA-dependent RNA polymerase (RdRp) (nsp12)
- Some accessory genes – 7-9

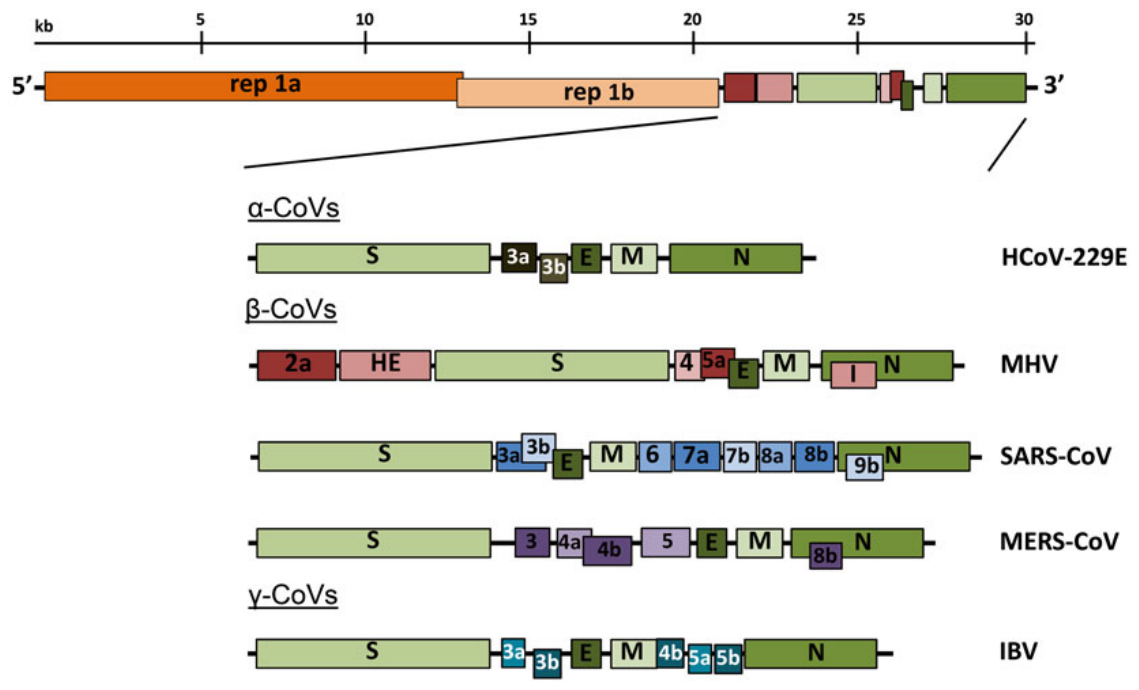

**Virion – General Coronaviruses**

- Round virus with club-like spikes – resembles solar corona (Latin for ‘crown’), hence the name
- Enveloped – lipid bilayer
- Spike protein (S)
  - 150 kDa
  - Homotrimer
  - Spike-like projection on surface of virion
  - Attaches to host cell receptor
  - Mediates fusion with host cell (class I fusion protein)
- Membrane protein (M)
  - Most abundant structural protein
  - 25-30 kDa
  - 3 transmembrane domains
  - Gives shape to virion – promote curvature?
  - Binds to nucleocapsid
- Envelope protein (E)
  - 8-12 kDa
  - Ion channel activity
  - Important for assembly and release of virus
- Nucleocapsid protein
  - Two domains that each bind RNA
  - Binds RNA like beads on a string
  - Binds to M
- Accessory proteins
  - Some function to block immune response

**Infection – Spike Protein – SARS-CoV2**

- 150 kDa protein that homotrimerizes
- Needs proteolytic cleavage to generate S1 and S2 in order to bind cells
- S proteins of coronaviruses are cleaved by a furin-like protease, but S protein of SARS-CoV2 is cleaved by furin itself. Furin is very abundant and widely expressed, so virus is easily activated
- S1 binds to the cell surface receptor protein, the receptor for SARS-CoV2 is ACE2 (angiotensin converting enzyme 2)
- S2 mediates fusion of virion and cell, buts needs to be proteolytically cleaved to expose the fusogenic peptide for fusion to occur
- Virions can internalize and S2 can be cleaved in late endosomes/lysosomes by cathepsin-like proteases
- S2 can be cleaved by TMPRSS2, a protease on the surface of the cell
- Fusion of virion membrane with cell/endosome/lysosome membrane releases nucleocapsid into cytoplasm

**Viral life cycle – General Coronaviruses**

**Fusion**

- In general, cathepsins, TMPRRS2 or other proteases can expose fusion peptide
- In some viruses, fusion takes place in acidified endosomes
- In some viruses, fusion takes place at the plasma membrane
- Fusion peptide on S2 inserts into cell membrane
- Heptad repeats in S2 interact to form 6-helix bundle
- Membranes fuse

**Translation – General Coronavirus**

- +ve stranded, single-stranded RNA is translated by ribosomes
- 2 ORFs, rep1a and rep1b, encode large polyprotein
- Rep1a and rep1b in different reading frames
- RNA pseudoknot stalls ribosome
  - ribosome unwinds knot and translates to rep1a stop codon

OR

- ribosome slips on slippery sequence to shift frame to translate rep1b
- polyprotein contains 16 nonstructural proteins (nsp)
- viral proteases cleave nsps (nsp3 and nsp5)
- RNA dependent RNA polymerase (RdRp) is nsp12
- Nsp7 and nsp8 – accessory proteins for RdRp
- Nsp13 is RNA helicase
- Nsp14 has exonuclease activity (ExoN) for proofreading
- Nsp1, nsp16 and nsp3 might block innate immune responses

**Replication and Transcription – General Coronavirus**

- Produces genomic and subgenomic RNAs. Latter translated to make structural proteins
- Requires a -ve stranded RNA intermediate to make genomic RNA and mRNAs
- Sequences in the 5’ untranslated sequence (5’-UTR) and 3’-UTR of the genome are important cis elements for replication
- Can undergo recombination – linked to the strand-switching ability of RdRp

**Assembly – General Coronavirus**

- After sub-genomic RNA synthesis, viral structural proteins are translated
- S, E and M are translated and inserted into ER
- Transit to ER-Golgi intermediate compartment (ERGIC), packaging occurs here
- Viral genome encapsidated by N protein bud into membranes with viral protein
- Mature virions released into lumen
- M, E and N needed to efficiently assemble virions
- S is incorporated last, before budding

**ACE2 bullets**

**Expression**

- Tissue level
  - Lung
  - Arteries
  - Heart
  - Kidney
  - Intestine
  - Testes
  - Thyroid
  - Adipose
  - Nasal and oral mucosa
  - Very low expression if any in
    - Spleen
    - Thymus
    - Lymph node
    - Bone marrow
    - Brain
    - Muscle
- Cellular level
  - Endothelial cells in arteries and veins in all tissues
  - Vascular smooth muscle cells
  - Myofibroblasts
  - Adipocytes
  - Type II pneumocytes in the lung
  - Type I pneumocytes in the lung
  - Enterocytes in small intestine – brush border staining
  - Basal layer of epidermis and smooth muscle in epidermis
  - In the brain – only see in endothelial cells and smooth muscle cells

**Function**

**Renin-angiotensin system**

Has systemic effects,

- Regulates blood pressure
- Fluid and electrolyte balance
- Systemic vascular resistance

but also local effects in organs and tissues that can be very important.

- E.g. regulation of learning and memory in the brain

**Regulation**

- Low blood flow in kidney, juxtaglomerular cells release renin into blood
- Renin cleaves angiotensinogen (from liver) to angiotensin I (AngI), a 10 amino acid peptide
- ACE cleaves AngI to AngII, an 8 amino acid peptide
- AngII signals via AT_1_R to mediate most of the effects associated with AngII, e.g. vasoconstriction
- AngII signals via AT_2_R to elicit the opposite effects, e.g. vasodilation

**ACE2**

- A carboxypeptidase
- Cleaves AngII to generate Ang(1-7)
- Ang(1-7) signals via Mas to elicit effects
- Effects oppose AngII effects
- Ang(1-7) effects
  - Vasodilation
  - Antithrombotic effect

**SARS-CoV2 Variants**

CDC classifies variants as ‘of interest’, ‘of concern’ or ‘of high consequence’

Variants of Interest:

- Mutation predicted to alter transmission, detection, treatment or resistance to immunity
- Causes increased number of cases
- Found in outbreak clusters

Variants of Concern, in addition:

- Widespread effect on detection via testing
- Substantially decreased response to therapy
- Significant decreased neutralization
- Reduced protection from vaccination
- Increased transmissibility
- Increased disease severity

Variants of High Consequence, in addition:

- Failure of diagnostics
- Significantly reduced vaccine effectiveness
- Significantly reduced susceptibilities to therapy
- More severe disease
- Increased risk of hospitalization
- **Note that there are no variants of high consequence at this time**

| **INFORMATION ON SARS-CoV2 ACCORDING TO THE CDC** | | | | | |
| --- | --- | --- | --- | --- | --- |
| **Strain** | **CDC Designation** | **Notable Spike Mutations** | **Transmission** | **Response to mAb** | **Neutralization*** |
| B.1.1.7 (Alpha) | variant of concern | E484K (some) | ~50% increase |  | minimal change |
| B.1.351(Beta) | variant of concern | K417N, E484K, N501Y | ~50% increase | significantly reduced | reduced |
| B.1.617.2 (Delta) | variant of concern | L452R | increased | potential reduction | potential reduction |
| P.1 (Gamma) | variant of concern | K417T, E484K, N501Y |  | significantly reduced | reduced |
| B.1.526 (Iota) | variant of interest | L452R, E484K (some) |  | reduced | reduced |
| B.1.427 (Epsilon) | variant of interest | L452R | ~20% increase | modest decrease | reduced |
| B.1.429 (Epsilon) | variant of interest | L452R | ~20% increase | modest decrease | reduced |
| B.1.617 (Kappa) | variant of interest | L452R |  | possible decrease? | possible reduction? |
| B.1.525 (Eta) | variant of interest | E484K |  | possible decrease? | possible reduction? |
| *Neutralization by convalescent or post-vaccination sera | | | | | |

Alpha variant has 69del and 70del. Deletions of these codons alters the spike gene sequence recognized by SARS-CoV2 tests.

**SESSION 4 -Coronavirus problem – pathogenicity**

*Recap – what were last week’s learning issues? What did we find out?*

***Discussion Topics***

- Any learning issues from last week that the students don’t address

*SARS-CoV2 infects the respiratory system. Describe the general organization of the respiratory system. Relate the general organization to the function of the respiratory system. Describe the cellular organization of the alveoli and how the cells relate to function.*

***Some Possible Discussion Topics***

- Why is the organization of the respiratory tract called a bronchial tree?
- Why do you breathe?
- What is the major function of the lungs?
- How do you think cells in the lungs are modified to facilitate their main function?
- How is oxygen delivered from the lungs to the body?
- How do you think lung cells and blood vessels are organized?

*People infected with SARS-CoV2 exhibit a wide range of symptoms and severity of disease. Some people are asymptomatic, others have mild symptoms, some have more severe symptoms and require hospitalization, while the most severely affected may die.*

*What symptoms are associated with SARS-CoV2 infection?*

***Some Possible Discussion Topics***

- What symptoms are associated with other viral infections, like the flu?
- How does the body respond to a viral infection?
- What symptoms might be associated with viral infection of the respiratory system?
- What symptoms associated with the cardiovasculature might be seen?
- What neurological/neurosensory symptoms might be seen?

*Co-morbidities are associated with the most severe cases of COVID-19. What are these co-morbidities?*

***Some Possible Discussion Topics***

- What are co-morbidities?
- What pre-existing conditions are associated with severe cases of COVID-19?
- Why might these comorbidities make people susceptible to viral infection?
- How might viral infection exacerbate comorbidities associated with COVID-19?

*What are the pathological effects in severe cases of COVID-19?*

***Some Possible Discussion Topics***

- What type of pathological damage would you expect by a respiratory virus?
- As the body responds to an infection, what might you see in infected tissue?
- How are immune cell responses controlled and what molecules might you detect in tissue and blood?
- Given the cardiovascular comorbidities, what pathology might you expect to see in the cardiovasculature?

**xxxxxxxxxxxxxxxxxxxxxxxxxxxxxxxxxxxxxxxxxxxxxxxxxxxxxxxxxxxxxxxxxxxxxxxxxxxxxxxxxxxx**

***Facilitator Cheat Sheet***

**Respiratory System**

Like a tree – trachea branches into smaller bronchi, branch into bronchioles. All these conduct air into the lungs and are lined with ciliated epithelium.

In the lungs are alveolar sacs. The alveoli have very thin walls of tissue where gas exchange takes place.

3 main types of cells:

- Type I alveolar cells (pneumocytes) – 95% of surface area
  - Very flattened
  - Gas exchange occurs here
  - Also provides barrier function
- Type II alveolar cells (pneumoctyes)
  - Blocky, cuboidal cells
  - Secrete surfactant
  - Divide to produce more Type I cells
- Capillary endothelial cells

Gas exchange occurs from air space across Type I cell, the basement membrane and the endothelial cell to the blood space

Images below from - <https://derangedphysiology.com/main/cicm-primary-exam/required-reading/respiratory-system/Chapter%20012/structure-and-function-alveolus>

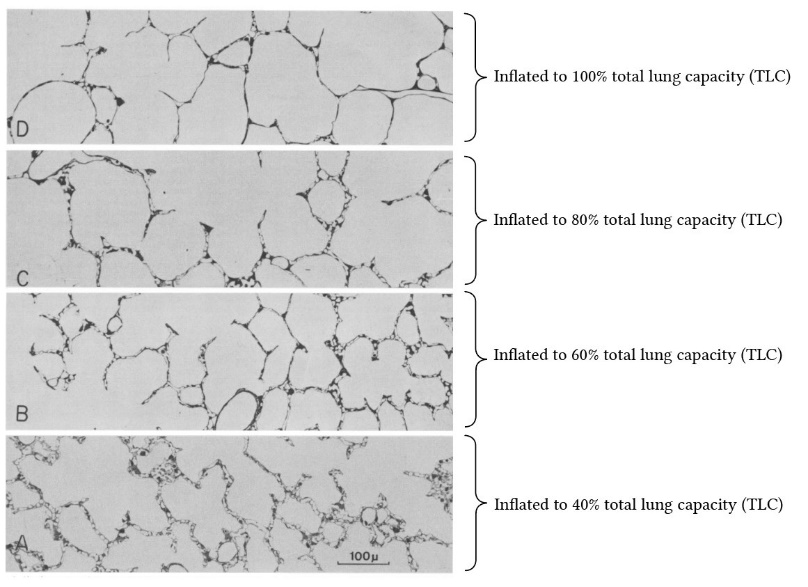

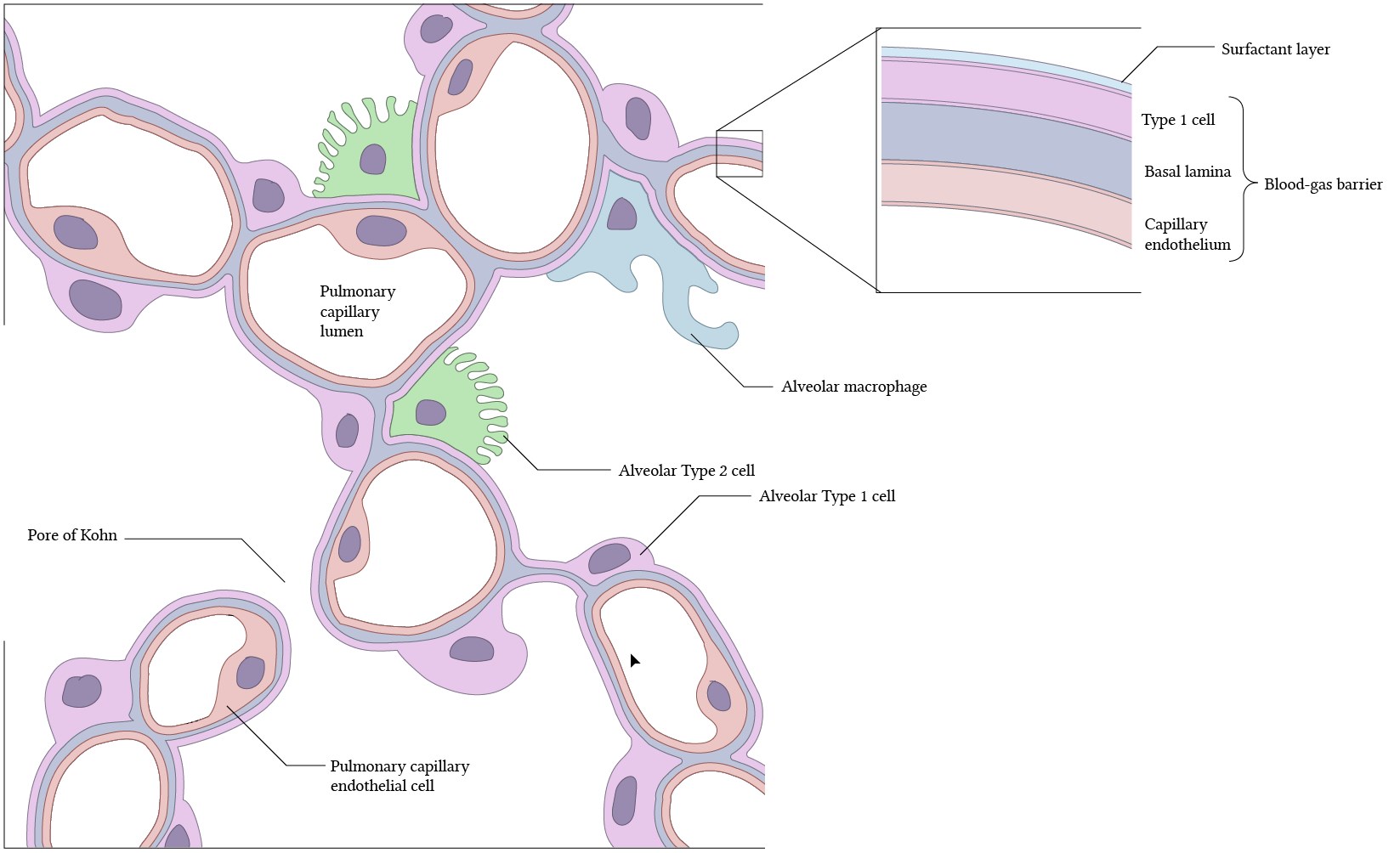

**Symptoms etc.**

**CDC says**

- **Incubation period**
  - Symptoms may appear 2-14 days after exposure
- **Symptoms**
  - Fever or chills
  - Cough
  - Shortness of breath or difficulty breathing (dyspnea)
  - Fatigue
  - Muscle or body ache (myalgia)
  - Headache
  - New loss of taste or smell – anosmia is loss of smell, ageusia is loss of taste
  - Sore throat
  - Congestion or runny nose
  - Nausea or vomiting
  - Diarrhea

**CT scan can reveal**

- Pneumonia
- Acute respiratory distress syndrome
- Acute cardiac injury

**Comorbidities**

**CDC – underlying medical conditions – Aug 2021 update**

<https://www.cdc.gov/coronavirus/2019-ncov/need-extra-precautions/people-with-medical-conditions.html>

Adults with these conditions can be at an increased risk of severe illness

- Chronic kidney disease
- COPD
- Immunocompromised state from organ transplant
- Obesity
- Serious heart conditions, heart failure, coronary heart disease or cardiomyopathies
- Sickle cell disease
- Type 2 diabetes
- Asthma
- Cerebrovascular disease or stroke
- Cystic fibrosis
- Hypertension or high blood pressure
- Immunocompromised state for other reasons
- Neurologic conditions, e.g. dementia
- Liver disease
- Pregnancy
- Pulmonary fibrosis
- Smoking
- Thalassemia
- Type 1 diabetes
- Cancer
- Down Syndrome
- HIV infection
- Transplant
- Substance abuse disorders

**Comorbidities from the literature**

| **Comorbidity** | **Richardson - NYC** | **Guan - China** | **Yang - China** |
| --- | --- | --- | --- |
|  | **n = 5700** | **n = 1590** | **n = 1567** |
| Hypertension | ~56.6% | ~16.9% | ~21.1% |
| Obesity | ~41.7% |  |  |
| Diabetes | ~33.8% | 8.2% | ~9.7% |
| Morbid obesity | ~19% |  |  |
| Coronary heart disease | ~11.1% |  |  |
| Asthma | ~9% |  |  |
| Congestive heart failure | ~6.9% |  |  |
| Cancer | ~6% |  |  |
| COPD | ~5.4% | ~1.5% |  |
| Chronic kidney disease | ~5% |  |  |
| End-stage kidney disease | ~3.5% |  |  |
| Sleep apnea | ~2.9% |  |  |
| Immunosuppression | ~1.8% | ~0.2% |  |
| Cirrhosis | ~0.4% |  |  |
| Hepatitis B | ~0.1% | ~1,8% |  |
| Hepatitis C | ~0.1% |  |  |
| Cardiovascular disease |  | ~3.7% | ~8.4% |
| Cerebrovascular disease |  | ~1.9% |  |
| Kidney disease |  | ~1.3% |  |
| Respiratory disease |  |  | ~1.5% |
| % no comorbidities | ~6.1% |  |  |
| % 1 comorbidity | ~6.3% |  |  |
| % >1 comorbidity | ~88% |  |  |

**Richardson JAMA 2020**

<https://jamanetwork.com/journals/jama/fullarticle/2765184>

**Guan European Respiratory J 2020**

<https://www.ncbi.nlm.nih.gov/pmc/articles/PMC7098485/pdf/ERJ-00547-2020.pdf>

**Yang Int J Inf Dis 2020**

<https://reader.elsevier.com/reader/sd/pii/S1201971220301363?token=BA1851FBAFC943B27058886008445B0D4DB347A328355801999AAB0A0F67282B5E10AA968EBB5E17FB35430C90130FC4>

**Cardiovascular problems**

- 1/5 patients have signs of heart injury
- blood vessel injury
- clots
- arrhythmia
- stroke
- heart attack
- myocarditis – inflammation of the heart – some patients have presented this way
- stress cardiomyopathy – release of catecholamines affects the heart – some patients have presented this way
- venous thromboembolic disease
- Interesting SARS study – 12 yr follow up on 25 patients - 68% have hyperlipidaemia, 44% have CVD abnormalities and 60% have glucose metabolism disorders

**Bloodwork can reveal**

- Lymphopenia – reduced lymphocytes
- Elevated leukocytes/neutrophilia – elevated neutrophils
- Thrombocytopenia – reduced platelets
- Increased plasma pro-inflammatory cytokines
  - IL2
  - Il7
  - IL10
  - G-CSF
  - IP10
  - MCP1
  - MIP1A
  - TNFα
  - CXCL10
  - IFNα
  - IFNγ
  - IL1β
  - IL12
  - IL18
  - IL33
  - TGFβ
  - CCL2
  - CCL3
  - XXL5
  - CXCL8
  - CXCL9
- High immune-inflammation index
- Prolonged prothrombin time
- High levels of IL6
- High levels of C-reactive protein
- High levels of D-dimer

**Autopsy can reveal**

- Diffuse alveolar damage in the lung
- Desquamation of pneumocytes
- Presence of hyaline membranes in alveoli – mixture of dead cells, surfactant and proteins deposited on alveolar walls
- Edema in lungs
- Hemorrhage in lungs
- Fibrin deposits
- Increase alveolar macrophages
- T cell infiltration in lung – CD4+ and CD8+
- Thrombosis – thrombi seen in lungs – can also be in liver, kidney, heart microvasculature
- CD61+ megakaryocytes in lung vessels – more than normal – and actively producing platelets
- Microangiopathy – thickened, weakened vessel walls – bleed, leak protein, slow flow
- Vessel growth via intussusceptive angiogenesis – dynamic process modifying microcirculation structure
- Cardiomegaly
- Right ventricle dilatation
- Lung pathology of COVID-19 similar to SARS, but thrombi and platelet aggregation is COVID-19 specific

**Can find the virus in**

- upper respiratory tract
- lungs
- Type II pneumocytes
- Phagosomes in macrophages
- Blood
- Endothelial cells

**SESSION 5 – cardiovascular and clotting problems associated with COVID-19**

*Recap – what were last week’s learning issues? What did we find out?*

***Discussion Topics***

- Any learning issues from last week that the students don’t address

*The pathogenicity of SARS-CoV2 extensively involves the cardiovascular system. Describe the structure of a blood vessel and the functions of the different cell types in a vessel.*

***Some Possible Discussion Topics***

- What are the different cell types found in a blood vessel?
- How are cells organized in a blood vessel?
- What are the major mechanisms of homeostasis in the cardiovascular system?
- What are the functions of the different cells in the vessel?

*A considerable proportion of COVID-19 patients exhibit abnormal coagulation, which could contribute to the severity of the disease and patient outcome. What is the evidence that abnormal coagulation occurs in COVID-19 patients (review of some things discussed in session 3)?*

***Some Possible Discussion Topics***

- What were the pathologies of SARS-CoV2 that were observed in the blood vessels of patients?

*Blood clotting is a mechanism of hemostasis, i.e. a mechanism to maintain the proper volume of blood. How is blood clotting normally regulated?*

***Some Possible Discussion Topics***

- How is a blood clot formed?/What is a blood clot made of?
- What proteins are important for blood clotting?
- How do platelets contribute to blood clotting?
- What enzymatic activities are associated with proteins involved in blood clotting?
- How do you turn blood clotting on?
- How do you turn blood clotting off?

***Writing Assignment - due Friday September 3, 2021 at 2 PM***

*This writing assignment is based upon the blood clotting sessions. Write a background to establish the basis of the hypothesis and explain the rationale for the proposed experiments. Succinctly state the hypothesis. Describe your experimental plan using multiple approaches to test the hypothesis. The plan should demonstrate rigor and include all of the essential controls to demonstrate the validity of the proposed experiments. You should also discuss your proposed analysis of the data, your expected results and an interpretation of your expected results. Your document should be between 1200 and 1400 words (excluding references). The rubric for written assignments will be used to grade the document. You will receive written feedback on the assignment and your writing skills will improve if you modify your document in response to the written feedback.*

**xxxxxxxxxxxxxxxxxxxxxxxxxxxxxxxxxxxxxxxxxxxxxxxxxxxxxxxxxxxxxxxxxxxxxxxxxxxxxxxxxxxx**

***Facilitator Cheat Sheet***

**Blood Vessels**

<https://edu.glogster.com/glog/cardiovascular-system-3262/2e5rk39bwfg?=glogpedia-source>

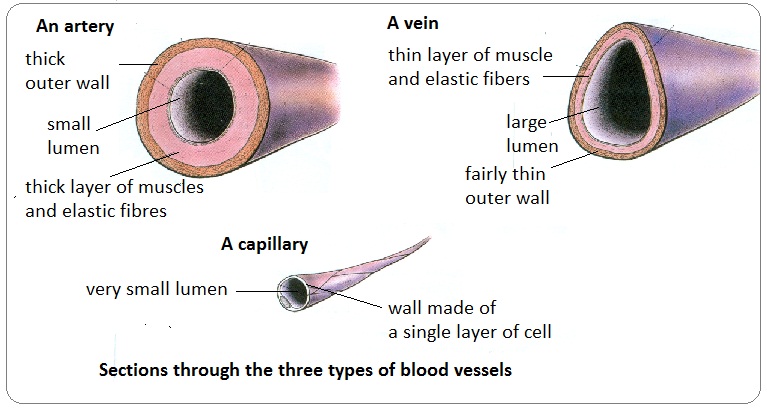

**Endothelial cells**

- Facilitate exchange of material between blood and tissue
- Normally anti-thrombogenic/anti-coagulant
  - 12-hydroxyoctadecadienoic acid (13-HODE)(13-OH-18:2) – chemorepellent
  - Produce prostacyclin
    - In response to platelet activation – e.g. ADP, thromboxane A_2_
    - Limit platelet aggregation
    - vasodilation
  - Produce NO
    - In response to platelet activation – e.g. ADP, thromboxane A_2_
    - Limits platelet aggregation
    - Vasodilation
    - Diminishes adhesion receptor expression on endothelial cells
  - Thrombomodulin – thrombin receptor on endothelial cell
    - Thrombomodulin-bound thrombin cleaves/activates protein C
    - Protein C inactivates V_a_ and VIII_a_ – i.e. is an anticoagulant
- Regulation of fibrinolysis
  - Releases tPA (tissue-type plasminogen activator) – constitutive and inducible
    - tPA cleaves and activates plasmin, which degrades fibrin, i.e. clots
  - Release plasminogen activator inhibitor type I (PAI-I) – constitutive and inducible
    - Inhibits tPA and uPA
- Can promote clotting
  - Synthesize and store von Willebrand factor
    - Stabilize factor VIII
    - Binds to tissue and platelets
  - Synthesize platelet-activating factor (PAF)
    - Activates platelets
    - Promotes platelet adhesion to endothelial cells
- Regulation of vascular tone
  - Prostacyclin and NO vasodilate
- Leukocyte binding and diapedesis
- Barrier function
  - Tight junctions and adherens junctions
  - Inflammation, sepsis, thrombin, cytokines, growth factors and ROS increases leakiness
    - Results in edema
- Regulate activity of circulating factors
  - E.g. ACE and ACE2 control activity of AngII

**Smooth muscle cells**

- Regulation of vascular tone
- Regulation of blood pressure

**From Session 3, pathogenicities linked to blood clotting**

- Clots
- Venous thromboembolic disease
- Thrombocytopenia
- Prolonged prothrombin time
- High levels of D-dimer
- Fibrin deposits
- Thrombi in lungs and other tissues
- Megakaryocytes in lung actively producing platelets

**Blood clotting**

- Rupture of vessel exposes underlying connective tissue
- Platelets adhere to connective tissue (collagen)
- Platelets activated – release factors to activate other platelets, e.g. ADP and thromboxane A_2_
- Activated platelets are sticky and bind fibrinogen, release ADP and thromboxane A_2_
- Platelets form a plug or soft clot
- Connective tissue contains Tissue Factor
  - Exposed Tissue Factor initiates activation of clotting factors
- Clotting factors mostly zymogens – inactive serine proteases activated by proteolytic cleavage
- One facter, e.g. VII_a_ cleaves an inactive zymogen, e.g. X, resulting in its activation, e.g. X_a_
- Different pathways can activate X_a_

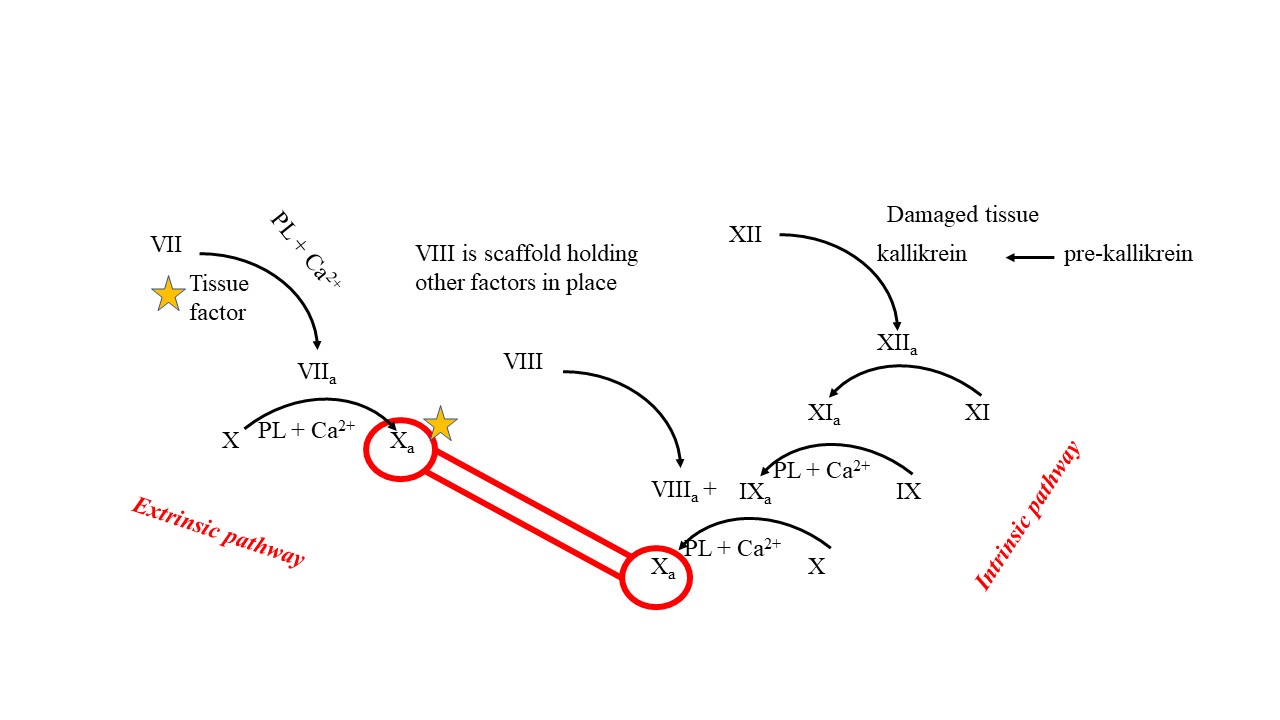

- X_a_ cleaves prothrombin (inactive) to thrombin (active protease)
- Thrombin cleaves fibrinogen (soluble in blood) to fibrin (can form insoluble fibers)
- Fibrin fibers form the stronger hard clot
  - Crosslinked to form a mesh like structure
- Platelets facilitate this cascade – clotting factors associate with the membrane of the platelets via calcium

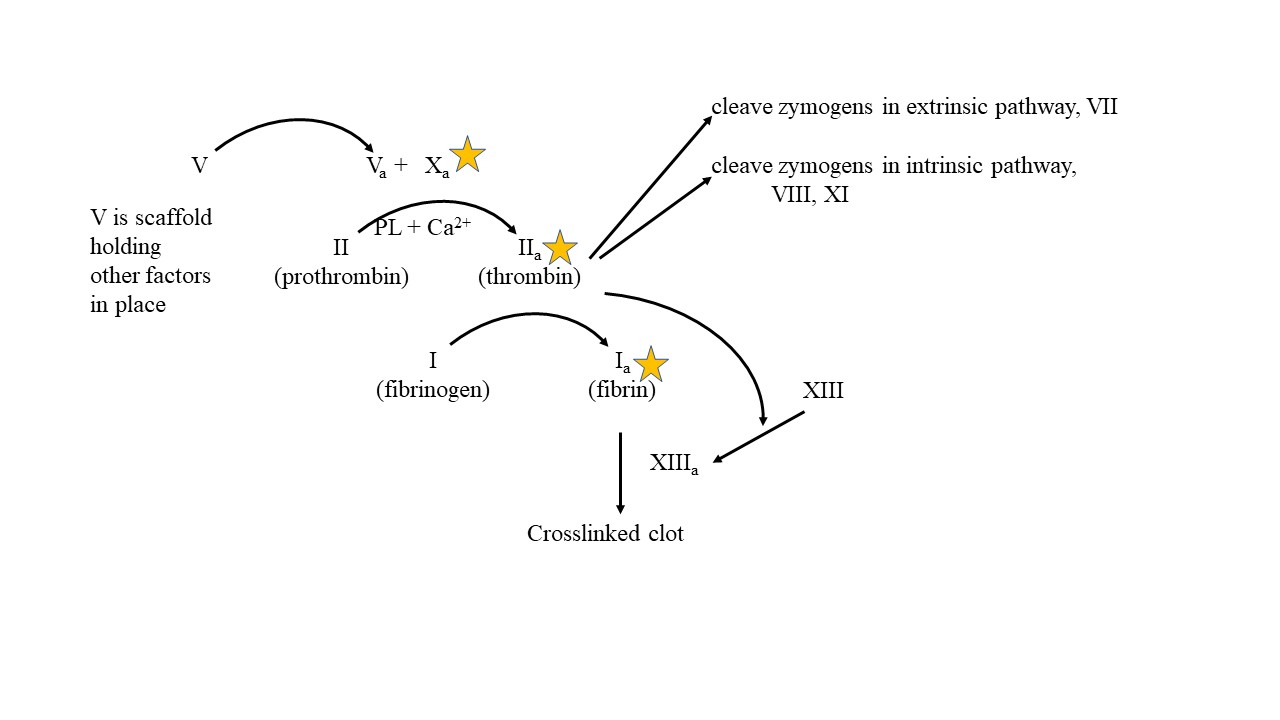

**Anticoagulation phase of clotting**

- Stop the cascade of clotting factors
- Protein C degrades V_a_ and VIII_a_
  - Activated by thrombin
- Serpins in blood (e.g. Antithrombin III (AT3)) – inhibits thrombin and X_a_
  - Heparin enhances effect

**Fibrinolysis**

- Remove the clot following wound healing
- tPA released from endothelial cells
  - cleave plasminogen to plasmin (active proteas)
- Plasmin cleaves fibrin
  - D-dimer is a product

**SESSION 6 – developing a hypothesis and beginning experimental design**

*Recap – what were last week’s learning issues? What did we find out?*

***Discussion Topics***

- Any learning issues from last week that the students don’t address

*We use the scientific method.*

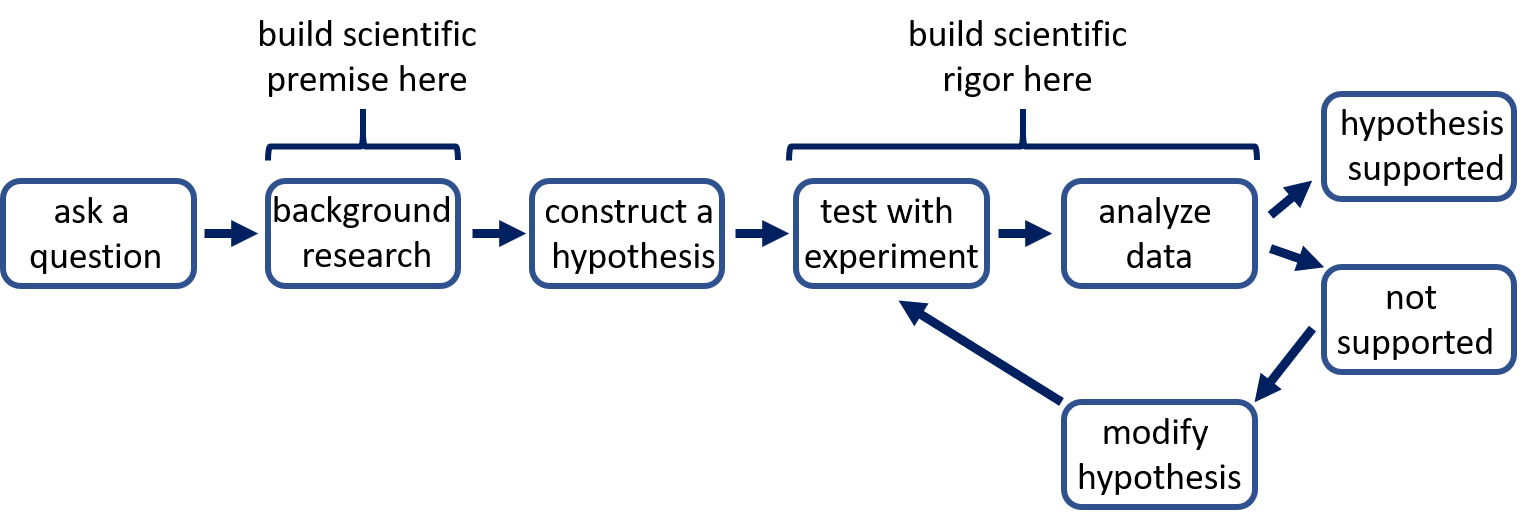

*A considerable proportion of COVID-19 patients exhibit abnormal coagulation, which could contribute to the severity of the disease and patient outcome.*

*An interesting and important question is:*

*What causes blood clotting in SARS-CoV2 infected people?*

*Do some research to learn about SARS-CoV2, the effects it has on infected individuals and how blood clotting is regulated. We have already discussed the SARS-CoV2 virus (Session 2), the pathogenesis of SARS-CoV2 (Session 3) and the regulation of blood clotting (Session 4).*

*What possible mechanisms might cause blood clotting in SARS-CoV2 infected individuals?*

**Some Possible Discussion Topics**

- How is blood clotting regulated?
- What pathogenicities are associated with SARS-CoV2?
- How are different systems altered by SARS-CoV2 infection?
- How is the immune system altered by SARS-CoV2 infection?
- How might pathologies/alterations of systems by SARS-CoV2 alter blood clotting?

*Consider the different possible mechanisms and develop one of these ideas into a hypothesis.*

*Begin the design of a series of experiments to test your hypothesis. What model system will you use? How will you manipulate the model system? What measurements will you make? What controls will be necessary?*

**Some Possible Discussion Topics**

- What type of model systems are possible to use to test the hypothesis?
- What problems can you foresee with each model system?
- What methods will you use to manipulate the system and what alternatives are available?
- What alternative approaches could be used to support the results of your experiment?
- What measurements could you make?
- What are all the controls you will need? Positive controls (does your system work)? Negative controls (is the outcome due to your manipulation)?

**xxxxxxxxxxxxxxxxxxxxxxxxxxxxxxxxxxxxxxxxxxxxxxxxxxxxxxxxxxxxxxxxxxxxxxxxxxxxxxxxxxxx**

***Facilitator Cheat Sheet***

**Model Systems**

- Mouse is a poor model system as mACE2 is different than hACE2 – virus doesn’t infect
  - Adenovirus vector used to express hACE2 in epithelial cells in respiratory system – intranasal administration
    - Infect and get high viral titer in lung
    - Get lung pathology and weight loss
  - Transgenic – K18 promoter drives hACE2 expression – epithelial expression in lung, liver, kidney and colon. Low mRNA levels in brain
    - Infect with SARS-CoV1
    - Weight loss, labored breathing
    - Lung path = hemorrhage, epithelial damage, congestion in alveolar system
    - Elevated inflammatory cytokines
  - Transgenic – CAG promoter, a composite promoter for high level expression in many tissues – express in spleen, stomach, heart, muscle, brain, kidney, lung, intestine and liver
    - Infect with SARS-CoV1
    - Lethargy, labored breathing, weight loss, death
    - Lung and brain affected
    - IP injection of virus – viremia, virus in brain, not in lung
  - HFH4-hACE2 transgenic – use a lung ciliated epithelial cell specific promoter – expresses in lung but also in brain, liver, kidney and GI
    - Infect with SARS-CoV1
    - Weight loss, death in 4-5 days
  - Transgenic – mACE2 promoter used to drive expression of hACE2 – express in lung, heart, kidney and intestine
    - Infect with SARS-CoV1
    - Pneumonia and extrapulmonary organ damage but mice didn’t die
- Macaques –
  - Rhesus macaques
    - Some weight loss, transient reduced appetite, increase respiration rate, hunched posture, change in piloerection, dehydration
    - Symptoms persist > 1wk
    - Find viral RNA in nose, pharynx, lung, gut and low levels in spinal cord, heart, skeletal muscle, bladder
    - Shed virus from nose and bronchioles/lung up to 5 dpi
    - Pathology = moderate interstitial pneumonia – thickened alveolar septa, edema, fibrin, hyaline membranes, increase alveolar macrophages, degeneration of epithelium, infiltration, ground-glass opacification,
    - Bloodwork – neutropenia, decreased hematocrit
    - Cytokine/chemokine levels up transiently
    - IHC show virus in type I and II pneumocytes, alveolar macrophages
  - Cynomolgus macaques
    - Infected animals – virus in lungs, trachea
    - Diffuse alveolar damage in lungs
    - Fluid in lungs
    - Virus in type I and II pneumocytes
    - Necropsy at 4 dpi – too early to see some phenotypes?
- Ferrets –
  - Elevated temp 2 dpi to 8 dpi
  - Reduced activity 2 dpi to 6 dpi
  - Occasional cough
  - No weight loss or death
  - See viral RNA in serum (low and at 2dpi and 4dpi)
  - See viral RNA and infectious virus in saliva and nasal wash 2dpi to 8 dpi.
  - See viral RNA in urine and feces
  - Necropsy and detect RNA in nose, trachea, kidney, lung, and GI. Only get infectious virus from nose and lung
  - IHC for viral antigens – in nose, trachea, lung and GI
  - RNA-seq for cytokines – up at 3 dpi and 7 dpi. 14 dpi IL6, IL1RN and IL1RA are still high
  - Path – immune infiltration, debris in alveolar wall
- Lung tissue explants
  - Can infect and produce virus
  - Induces IFN and cytokines/chemokines
- Cells

**Manipulations**

- Genetics – KO, Transgene, CRISPR, shRNA
- Small molecule inhibitors
- Antibodies

**Measurements**

- Full blood count - platelet count is part of this
- Bleeding time test
- Activated partial thromboblastin time (aPTT)
  - External agent used to activate factor XII, e.g. kaolin, a type of clay can do this (activates top of the extrinsic pathway)
- Prothrombin time test (PT)
  - Use tissue factor to activate factor VII (activates top of the intrinsic pathway)
- Thrombin time (TT)
  - Add thrombin to initiate clotting (activates down in the common pathway)
- Levels of D-dimer
- Fibrin deposits
- Thrombosis
- Megakaryocytes producing platelets in vessels
- Direct measure of amounts of clotting factors in the blood
- Platelet aggregation test

**SESSION 7 – building experimental design**

*Recap – what were last week’s learning issues? What did we find out?*

***Discussion Topics***

- Any learning issues from last week that the students don’t address

*Review your model system and experimental design, including methods, measurements and controls. Are the methods and experimental approach rigorous? How could they be made more rigorous?*

***Discussion Topics***

- How many replicates or how many animals do you need for the study?
- What other methods and measures would support your experiment?

*Discuss how your controls validate the outcome of the experiment. What additional controls could increase confidence in your result?*

***Discussion Topics***

- What other positive controls could be used?
- What other negative controls could be used?

*How are you going to analyze your experimental results?*

***Discussion Topics***

- How are you going to look at your data and present your data?
- What analyses can you do to determine if the results of the controls and the experimental are truly different?

*What are your expected results and how will you interpret these results?*

***Discussion Topics***

- What do you expect to see in your measurement if the hypothesis is correct? Incorrect?
- What other explanations can result in getting the measurement you expect, yet your hypothesis is incorrect? What control might rule this out.

*If your hypothesis is supported by the results of your experiments, what will you do next?*

*If your hypothesis is not supported by your results, what modifications to your hypothesis or alternative hypothesis will you propose? How would you test this (superficial description)*

**SESSION 8 – neurological problems associated with COVID-19, particularly anosmia and ageusia**

*The pathogenicity of SARS-CoV2 involves the nervous system. Generally review the symptoms/pathogenicities associated with the nervous system.*

***Some Possible Discussion Topics***

- What are the symptoms with which patients might present?
- What neurological/neurosensory symptoms might be seen?
- What pathogenicities are associated with COVID-19?
- Which of these might be related to the nervous system?

*A considerable proportion of COVID-19 patients exhibit anosmia (loss of smell). Describe the structures required for the sense of smell and the processes required to transmit the presence of an odor molecule in the nose to a signal in the brain.* *How is this altered in some people infected with SARS-CoV-2?*

***Some Possible Discussion Topics***

- What different cell types are in the olfactory epithelium?
- How are signals from the olfactory epithelium transmitted to the brain?
- What is the molecular mechanism for detection of odors?
- How is odor detection translated into an action potential?
- What happens when olfactory sensory neurons die?
- What effect does SARS-CoV-2 infection have on the olfactory epithelium?
- Does the virus infect olfactory sensory neurons?

*A considerable number of COVID-19 patients exhibit ageusia (loss of taste). Describe the structures in the tongue that are responsible for taste and the processes required to transmit the presence of a taste molecule on the tongue to a signal in the brain. How is this altered in some people infected with SARS-CoV-2?*

***Some Possible Discussion Topics***

- What different cell types are in the taste buds?
- How are signals from the taste buds transmitted to the brain?
- What are the molecular mechanisms for detection of tastes?
- How is taste detection translated into an action potential in different taste bud cells?
- What happens when taste bud cells die?
- What effect does SARS-CoV-2 infection have on taste buds?
- Does the virus infect cells in the taste buds?

**xxxxxxxxxxxxxxxxxxxxxxxxxxxxxxxxxxxxxxxxxxxxxxxxxxxxxxxxxxxxxxxxxxxxxxxxxxxxxxxxxxxx**

***Facilitator Cheat Sheet***

**COVID-19 and the nervous system**

**Symptoms**

- Headache
- Confusion
- Dizziness
- Anosmia
- Ageusia
- Seizure
- Numbness

**Pathogenicity**

- Cerebral hemorrhage
- Cerebrovascular disease
- Encephalitis
- Edema
- Vasculitis
- Guillain- Barré Syndrome
- Stroke

**Presence of virus**

- Can detect in CSF
- EM shows virus in brain and endothelial cells at BBB
- In astrocytes and microglia
- Some say you can see in neurons

**Modes of transmission**

- Blood
- Retrograde transport via olfactory network

**Mechanisms of neurological effects**

- Presence in these tissues suggests direct effects could occur
- Indirect due to cytokine storm
- Hypoxemia

**The sense of smell**

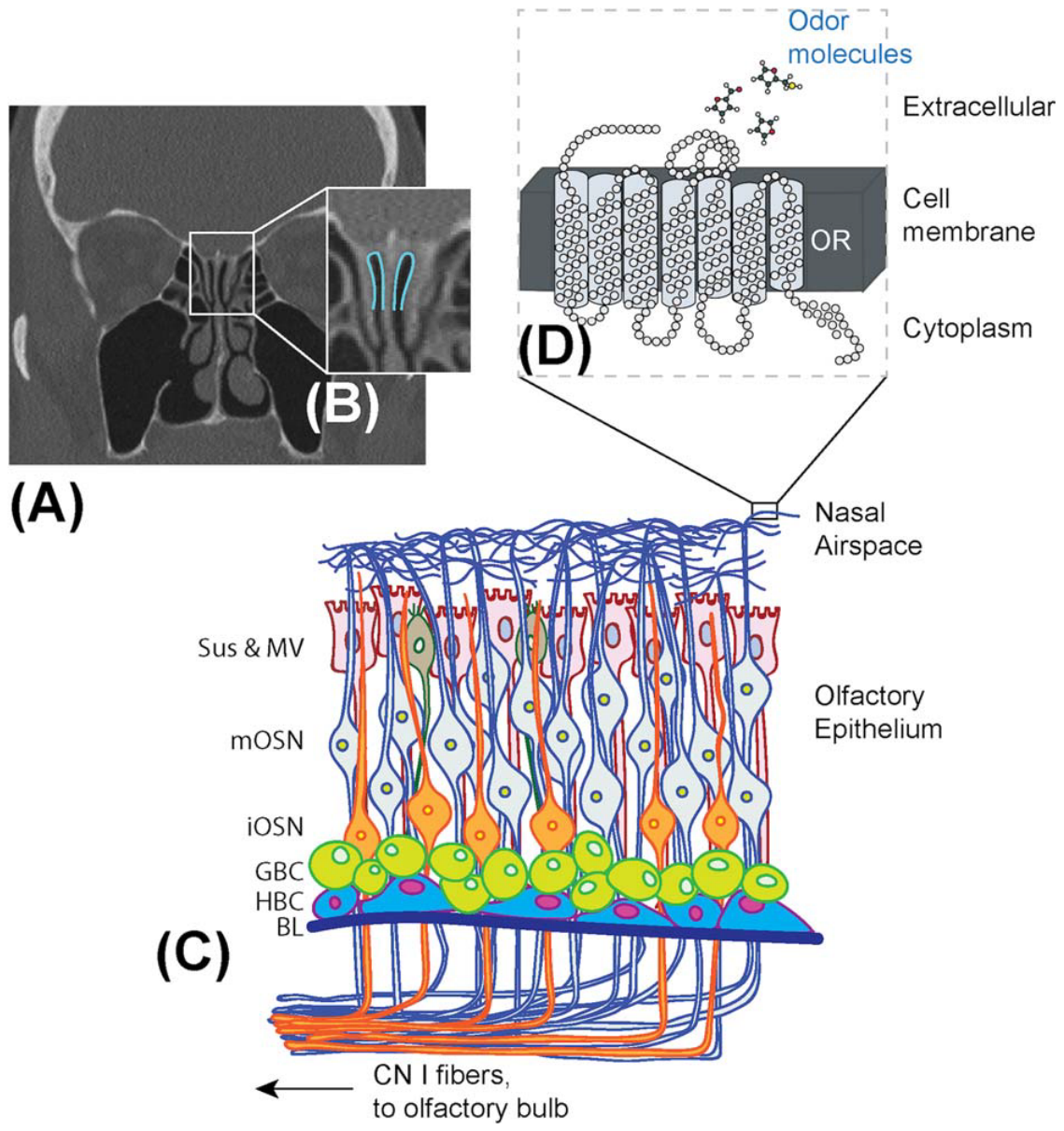

**From Choi and Goldstein, 2018 Laryngoscope Investigative Otolaryngology 3:35-42.**

**Olfactory Epithelium (OE)**

- Pseudostratified
- Multiple cells types
  - Olfactory sensory neurons (OSN)
    - Mature OSN
    - Immature OSN
  - Stem cells – horizontal basal cells
    - Renewal of OE, including neurons, as needed
  - Progenitor cells – globose basal cells
    - Renewal of OE, including neurons, as needed
  - Sustentacular cells (glia like)
  - Sensory microvillar cells
- Need air conduction to get smells to the OE, which is high up in the nose

**Olfactory Sensory Neurons (OSN)**

- Dendrites extend to surface of OE
- Immotile cilia extend from dendrites into the air space
- Cilia have receptors for smells - GPCRs
- Axon projects from cell body and runs to the olfactory bulb
- Axons bundled to make olfactory nerve (CN I)
- In olfactory bulb axons synapses of mitral and tufted cells in structures called glomeruli
- OSN only live for a few months and then need to be replaced by horizontal basal cells and globose basal cells

**Odor Signal Transduction**

- Receptors are GPCRs – about 700 different olfactory GPCRs in humans
- Detect odorants that are <300 MW and hydrophobic, maybe have an aldehyde
- Each OSN expresses only 1 type of GPCR – sensitive to 1 GPCR
- GPCRs are coupled to G_αolf_
- Signal goes to adenylyl cyclase and increase cAMP
- cAMP opens cyclic nucleotide gated channel causing depolarization
- depolarization opens voltage gated channels in axon hillock to trigger action potential along the axon
- neurotransmitter release at synapse in olfactory bulb

**Mechanisms of anosmia**

- Infections with a number of different viruses can cause degeneration of OE – cause of anosmia
- Obstruction of airflow can cause anosmia
- Rhinosinusitis – inflammation – can cause anosmia
- Inflammatory cytokines can directly affect OE cells
- Cytokine exposure can cause OSN death
- Inflammation can impair OE basal cell proliferation

**The sense of taste**

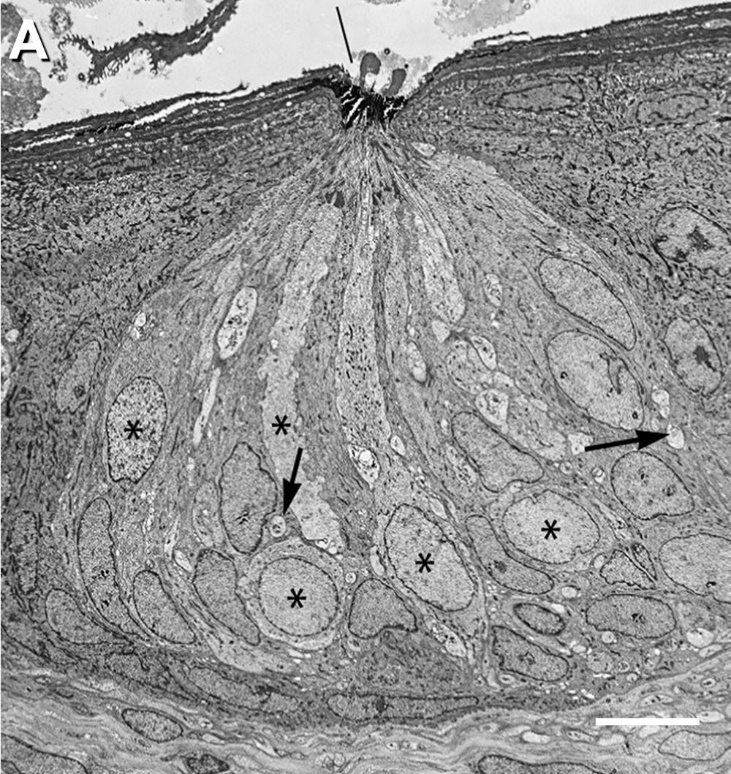

**Taste bud from Chaudhari and Roper, 2010, J Cell Biol 190:285-296**

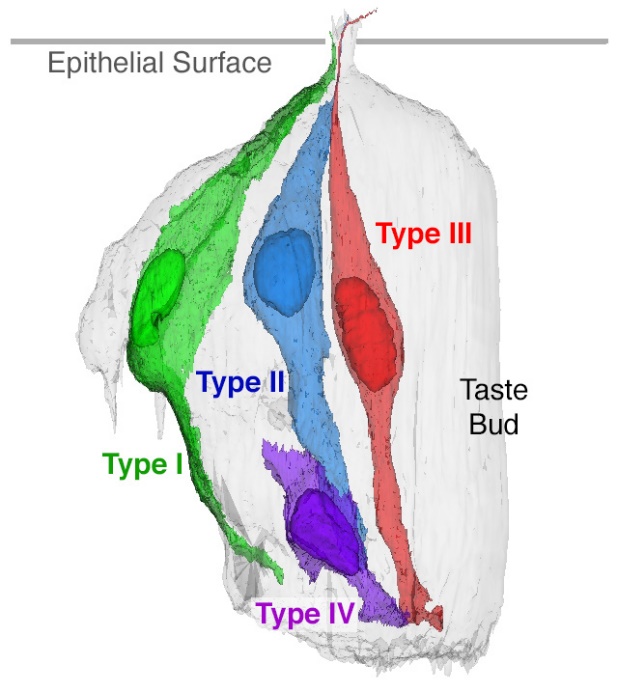

**From** Kinnamon SC and Finger TE. **Recent advances in taste transduction and signaling [version 1; peer review: 2 approved]** F1000Research 2019, **8**(F1000 Faculty Rev):2117 (<https://doi.org/10.12688/f1000research.21099.1>)

**Lingual Papillae**

- Structural projections on the tongue
- Filiform papillae – function in sensing touch – have no taste buds
- Fungiform, foliate and circumvallate papillae all have taste buds

**Taste buds**

- Clusters of neuroepithelial cells (modified epithelial cells) – 50 - 100 cells
- Four cell types, Type I, Type II, Type III and Type IV. Type I-III are differentiated, Type IV is not
- Avg lifespan of taste cells is 10 – 14 days
  - Old cells undergo apoptosis
  - New cells enter the taste bud (become Type IV cells), which differentiate in a couple of days
- Progenitor cells are Keratin 5 +ve and Keratin 14 +ve. These are in epithelium outside taste buds
  - Sonic Hedgehog +ve cells differentiate into Type I, Type II and Type III cells
  - Sonic Hedgehog -ve cells differentiate into non-taste epithelium of the tongue
    - These turnover every 5 days

**Type I cells**

- Performs functions that glia cells do in other places
- Most abundant, wrap lamellae around other cells
- Might function to remove neurotransmitters
  - Express GLAST – glutamate transporter
  - Express cell surface NTPDase2 – hydrolyzes extracellular ATP
  - Express ROMK – a K+ channel – removes extracellular K+

**Type II cells**

- Has receptors for sweet, bitter and umami – these are GPCRs – many different GPCRs
- A cell only expresses one type of GPCR
- No prominent synapses, nerve fibers do come close to these cells
- Unconventional neurotransmission mechanism?

**Type III cells**

- Forms synapses
- Makes GABA and 5-HT (serotonin)
- Respond to salt, sour and carbonated solutions

**Type IV cells**

- Basal, round cells

**Signal transduction in Type II cells**

- Different sets of GPCRs for bitter, sweet and umami
- Main signal is via βγ activation of PLCβ2
- PLCβ2 generates IP3 and Ca2+ dump.
- Ca2+ binds cation channel TRPM5 and gap junction hemichannel
- TRPM5 opening generates action potential via voltage gated Na+ channels
- Gap junction hemichannels – voltage gated large pore channel with two subunits CALHM1 and CALHM2 – ATP and other molecules pass out of the cell
- Need cytosolic calcium and depolarization for ATP release
- ATP release stimulates afferent nerve fibers directly and Type III cells, which then release 5-HT and NE
- ATP also acts in autocrine fashion on type II cells – positive feedback to release

**Signal transduction in Type III cells**

- Sour taste
  - OTOP1 – an ion channel, is opened allowing H+ to enter
  - Triggers depolarization directly and by blocking K+ channels
  - Depolarization triggers voltage gate Na+ channels
  - Generates action potentials to activate voltage gated Ca2+ channels causing synaptic vesicle release
- Salt taste
- Two paths –amiloride-sensitive (AS) and amiloride-insensitive (AI) mechanisms
- AS is medicated via ENaC
- High concs of salt thought to activate a subset of Type II and a subset of Type III cells

**Nerve fibers**

- 3-14 sensory neurons connect to each taste bud – depends on species and area of mouth
- Some form synapse with Type III cells, others just wind around the cells in the taste bud
- Cell bodies of these sensory neurons in geniculate, petrosal and nodose cranial ganglia, in CN VII, CN IX and CN X.
- In absence of innervation, taste buds degenerate
- A population of the ganglion cells express a touch sensitive marker – maybe involved in the texture part of taste.

**Mechanisms of ageusia**

- Innate immunity via IFN and TNFα – bystander effect
- Innervation and degeneration of taste buds
- IFN can induce ACE2 expression

**COVID-19 and the loss of sense of smell and taste**

CDC says new loss of taste or smell is symptom seen 2-14 days after exposure

- Frequency varies in individual studies – large compilation of studies suggests 75-80% or more of the patients report anosmia or ageusia
- A majority of patients recover

ACE2 is widely expressed in oral epithelium and in nasal epithelial cells

- In the olfactory epithelium, ACE is expressed in non-neuronal cells - studies say neuronal cells are ACE2 -ve
- TMPRSS2 is expressed in neuronal and non-neuronal cells
- ACE2 expression in oral cavity
  - Squamous epithelium
    - ACE2 in basal and spinous layers
    - TMPRSS2 in all layers
  - Taste buds
    - ACE2 expressed in epithelial layer
    - TMPRSS2 expressed in taste buds
  - Submandibular glands
    - Serous cells express ACE2 and TMPRSS2
    - Ductal epithelium expresses ACE2 and TMPRSS2
  - Single cell RNA seq data – Cell clusters with high ACE2 expression have very low levels of markers of taste bud cells, markers of taste cell progenitors and basal epithelial cells. Very low percentage of taste bud cells express ACE2.

**SESSION 9 – developing a hypothesis and beginning experimental design**

*Recap – what were last week’s learning issues? What did we find out?*

***Discussion Topics***

- Any learning issues from last week that the students don’t address

*A considerable proportion of COVID-19 patients exhibit anosmia and ageusia.*

*“The Strange Grief of Losing My Sense of Taste.*

*Many symptoms of Covid-19 were difficult, but losing by ability to taste hurt the most.”*

*OpEd in the New York Times, Sept 6, 2020 by Krista Diamond*

*An interesting and important question is:*

*What causes the transient ageusia in SARS-CoV-2 infected people?*

*Do some research to learn about SARS-CoV2, the effects it has on infected individuals and how the sense of taste operates. We have already discussed the SARS-CoV2 virus (Session 2), the pathogenesis of SARS-CoV2 (Session 3) and the senses of smell and taste (Session 7), i.e. we did this in advance.*

*What possible mechanisms might cause ageusia in SARS-CoV2 infected individuals?*

**Some Possible Discussion Topics**

- What structures/cells are responsible for sensing taste?
- What cells express ACE2 and TPMRSS2?
- How are chemical signals in the oral cavity transmitted via sensory neurons?
- What are the physiological responses to viral infection?

*Consider the different possible mechanisms and develop one of these ideas into a hypothesis.*

*Begin the design of a series of experiments to test your hypothesis. What model system will you use? How will you manipulate the model system? What measurements will you make? What controls will be necessary?*

**Some Possible Discussion Topics**

- What type of model systems are possible to use to test the hypothesis?
- What problems can you foresee with each model system?
- What methods will you use to manipulate the system and what alternatives are available?
- What alternative approaches could be used to support the results of your experiment?
- What measurements could you make?
- What are all the controls you will need? Positive controls (does your system work)? Negative controls (is the outcome due to your manipulation)?

**xxxxxxxxxxxxxxxxxxxxxxxxxxxxxxxxxxxxxxxxxxxxxxxxxxxxxxxxxxxxxxxxxxxxxxxxxxxxxxxxxxxx**

***Facilitator Cheat Sheet***

**Model Systems – animal models (from Session 5)**

- Mouse is a poor model system as mACE2 is different than hACE2 – virus doesn’t infect
  - Adenovirus vector used to express hACE2 in epithelial cells in respiratory system – intranasal administration
    - Infect and get high viral titer in lung
    - Get lung pathology and weight loss
  - Transgenic – K18 promoter drives hACE2 expression – epithelial expression in lung, liver, kidney and colon. Low mRNA levels in brain
    - Infect with SARS-CoV1
    - Weight loss, labored breathing
    - Lung path = hemorrhage, epithelial damage, congestion in alveolar system
    - Elevated inflammatory cytokines
  - Transgenic – CAG promoter, a composite promoter for high level expression in many tissues – express in spleen, stomach, heart, muscle, brain, kidney, lung, intestine and liver
    - Infect with SARS-CoV1
    - Lethargy, labored breathing, weight loss, death
    - Lung and brain affected
    - IP injection of virus – viremia, virus in brain, not in lung
  - HFH4-hACE2 transgenic – use a lung ciliated epithelial cell specific promoter – expresses in lung but also in brain, liver, kidney and GI
    - Infect with SARS-CoV1
    - Weight loss, death in 4-5 days
  - Transgenic – mACE2 promoter used to drive expression of hACE2 – express in lung, heart, kidney and intestine
    - Infect with SARS-CoV1
    - Pneumonia and extrapulmonary organ damage but mice didn’t die
- Macaques –
  - Rhesus macaques
    - Some weight loss, transient reduced appetite, increase respiration rate, hunched posture, change in piloerection, dehydration
    - Symptoms persist > 1wk
    - Find viral RNA in nose, pharynx, lung, gut and low levels in spinal cord, heart, skeletal muscle, bladder
    - Shed virus from nose and bronchioles/lung up to 5 dpi
    - Pathology = moderate interstitial pneumonia – thickened alveolar septa, edema, fibrin, hyaline membranes, increase alveolar macrophages, degeneration of epithelium, infiltration, ground-glass opacification,
    - Bloodwork – neutropenia, decreased hematocrit
    - Cytokine/chemokine levels up transiently
    - IHC show virus in type I and II pneumocytes, alveolar macrophages
  - Cynomolgus macaques
    - Infected animals – virus in lungs, trachea
    - Diffuse alveolar damage in lungs
    - Fluid in lungs
    - Virus in type I and II pneumocytes
    - Necropsy at 4 dpi – too early to see some phenotypes?
- Ferrets –
  - Elevated temp 2 dpi to 8 dpi
  - Reduced activity 2 dpi to 6 dpi
  - Occasional cough
  - No weight loss or death
  - See viral RNA in serum (low and at 2dpi and 4dpi)
  - See viral RNA and infectious virus in saliva and nasal wash 2dpi to 8 dpi.
  - See viral RNA in urine and feces
  - Necropsy and detect RNA in nose, trachea, kidney, lung, and GI. Only get infectious virus from nose and lung
  - IHC for viral antigens – in nose, trachea, lung and GI
  - RNA-seq for cytokines – up at 3 dpi and 7 dpi. 14 dpi IL6, IL1RN and IL1RA are still high
  - Path – immune infiltration, debris in alveolar wall

**Model Systems - other**

- Organoids – can develop organoids in culture with full differentiation into taste cells
  - Taste cells can be stimulated with “tastes” and signal can be measured by calcium flux

**Measurements**

- Animal response to taste – lick tests
  - 2 bottle taste test – 1 bottle is water, other has 1 taste (e.g. sweet). Measure water drank out of each bottle over 96 hours. Calculate preference as a percentage of total water intake
  - Brief access test – provide water for 10 s, then tasty solution for 10 s. Measure number of licks of each over the 10 s period
  - Brief access taste aversion assay (BATA) – use to assess taste aversion of pharmaceuticals – One-week standardization protocol – Day 1 withdraw water, Day 2 restore water (training), Day 3 test but only use water, Day 4 test with the controls and compound, Day 5 test with the controls and compound – measures the number of licks. There are known compounds that are taste aversive to rats.
- Conditional taste aversion
  - Condition mice for taste aversion – give novel taste – inject LiCl IP to produce GI malaise – recover a day. Measure taste aversion with one-bottle or two-bottle test. Measure amount drank
  - This is a measure of taste and memory, so more complex
- Neuroimaging – is Tg mouse with GCaMP3 transgene expressed in sensory neurons. This is GFP modified calcium sensor. Need to drill a hole to image the geniculate ganglion.
  - Perfuse stimulus into mouth for 5 sec. Neurons in geniculate ganglion will light up. Repeat stimuli multiple times to see a pattern of neurons responding.
  - Can stimulate tongue with electric pulse to activate taste receptors and primary sensory afferent fibers. Triggers response in ganglion cells – control to make sure the effect is on the tongue and not a neuronal transmission defect
  - Can introduce GCaMP3 into any mouse using AAV vector, injection into specific places in the brain and retrograde transport to the cell bodies in the ganglia.

**SESSION 10 – building experimental design**

*Recap – what were last week’s learning issues? What did we find out?*

***Discussion Topics***

- Any learning issues from last week that the students don’t address

*Review your model system and experimental design, including methods, measurements and controls. Are the methods and experimental approach rigorous? How could they be made more rigorous?*

***Some Discussion Topics***

- How many replicates or how many animals do you need for the study?
- What other methods and measures would support your experiment?

*Discuss how your controls validate the outcome of the experiment. What additional controls could increase confidence in your result?*

***Some Discussion Topics***

- What other positive controls could be used?
- What other negative controls could be used?

*How are you going to analyze your experimental results?*

***Some Discussion Topics***

- How are you going to look at your data and present your data?
- What analyses can you do to determine if the results of the controls and the experimental are truly different?

*What are your expected results and how will you interpret these results?*

***Some Discussion Topics***

- What do you expect to see in your measurement if the hypothesis is correct? Incorrect?
- What other explanations can result in getting the measurement you expect, yet your hypothesis is incorrect? What control might rule this out.

*If your hypothesis is supported by the results of your experiments, what will you do next?*

*If your hypothesis is not supported by your results, what modifications to your hypothesis or alternative hypothesis will you propose? How would you test this (superficial description)?*

**SESSION 11 – SARS-CoV2 Therapeutics**

*What COVID drugs have been developed? What kind of drugs are they (small molecule or biological)? What is their target? What is their mechanism of action?*

***Some Possible Discussion Topics***

- What type of drugs might block infection of cells?
- What type of drugs might block the viral replication cycle?
- What type of drugs work on the virus or viral proteins?
- What type of drugs work on human proteins that are required for viral replication?

*Review the SARS-CoV-2 virus and lifecycle again, with a focus upon identifying additional targets for therapeutic intervention (Discussion in class).*

***Some Possible Discussion Topics***

- What proteins are found in the virus?
- What are the functions of these proteins (particularly the spike protein, S)?
- How does the virus attach to the cell – what is the receptor on the virus, what is the receptor on the cell?
- How is the receptor on the virus (S) modified to promote its function of cell attachment – what enzyme is responsible?
- What viral protein is responsible for promoting viral fusion?
- How are viral mRNAs and new viral genomes made?
- What viral proteins are required for transcription and replication?

*Select a novel target or a known target and consider how you would like new drugs to inhibit your target (mechanism). Design experiments for a high throughput assay to identify drugs against a novel target (“first in class”) or better drugs against known targets (“best in class”). The high throughput assay will require automation and the experiment needs to be as simple as possible (Discussion in class). Positive and negative controls* will be required for your assay.

***Some Possible Discussion Topics***

- Are you trying to target a protein-protein interaction or the function of an enzyme?
- How can you measure a protein-protein interaction in a very simple assay (what does your target protein bind to?)?
- What could you use as a readout for binding?
- How can you measure the activity of your enzyme in a very simple assay (what is the substrate of your target enzyme?)?
- What could you use as a reading for enzymatic activity?

*One learning issue for Session 11 – You have designed four variations on a screening assay for your drug. You run a pilot experiment with all four assays. The data is present in the excel spread sheet. Look at the data and pick which assay looks like the best to you. Then read the Z’ primer and calculate the Z’ for each assay. Did you pick out the best assay without doing the calculation? Which assay should you use moving forward?*

**xxxxxxxxxxxxxxxxxxxxxxxxxxxxxxxxxxxxxxxxxxxxxxxxxxxxxxxxxxxxxxxxxxxxxxxxxxxxxxxxxxxx**

***Facilitator Cheat Sheet***

**COVID-19 Drugs**

CDC currently lists the 4 highlighted drugs as treatments for outpatients (Aug 2022)

- REGN-COV2 – cocktail of neutralizing antibodies – block binding to cells. (FDA approved for emergency use)
- Remdesivir – adenosine analogue – RdRp inhibitor (FDA approved for emergency use – Oct 2020)
- Favipiravir – purine nucleotide analogue – RdRp inhibitor
- Plitidepsin and Zotatifin – inhibit cell translation factors to impair translation and viral replication (in trials)
- Bamlanivimab and Etesevimab – monoclonal antibodies targeting Spike protein blocking infection (FDA approved for emergency use)
- Molnupiravir – nucleoside analogue (cytosine) – RdRp inhibitor (FDA approved for emergency use – Dec 2021)
- Paxlovid (Nirmatrelvir and Ritonavir) – nirmatrelvir is 3C-like protease inhibitor (3C are class of proteases in coronaviruses – responsible for cleaving the polyprotein). Ritonavir is a protease inhibitor originally developed for HIV (FDA approved for emergency use – Dec 2021)
- Bebtelovmab – monoclonal antibody targeting spike protein to block infection – recognizes omicron (FDA approved for emergency use – Feb 2022)

**Drugs repurposed for COVID-19**

- Chloroquine and hydroxychloroquine – block endosomal acidification, inhibit glycosylation of receptor and proteolytic processing. Originally malaria drugs. (FDA approved for emergency use April 2020 and then unapproved in June 2020 – cardiovascular complications)
- Lopinavir/ritonavir (antivirals) – aspartate protease inhibitor/P450 inhibitor (latter increases lopinavir half-life in blood. Originally HIV drugs. (Trial ended by WHO in July 2020 – no mortality decrease)
- Ribavirin – guanine analogue – inhibits RdRp
- Umifenovir – kinda unclear – inhibits viral entry – binds flu HA preventing fusion

**Targets with some experimental evidence**

- Small molecules/drugs that bind S
- Peptides vs S2 also developed
- ACE2 inhibitor might also work if it blocks S binding
- Protease inhibitors – TMPRSS2, furin and cathepsin inhibitors could work – caveat might need broad spectrum inhibitors

**Additional Targets**

- Viral proteases that cleave the polyprotein (Nsp3 and Nsp5)
- Nsp12 – RdRp – target of remdesivir
- Nsp13 – a helicase – pretty important for replication – potential target
- Nsp14 – a 3’-5’ exonuclease – some proof reading function – potential target
- Nsp15 – uridine specific endoribonuclease - ??
- Nsp16 – RNA-cap methyltransferase -??

**Strategies for a drug screen**

Keep it simple as we are aiming for a high throughput screen. Simple binding assay or simple enzyme assay with a read out that is really simple, e.g. fluorescent assay

**Z’ Primer – for the Z’ learning issue**

Z’ is a statistic used to determine if an assay is suitable for high throughput screening. The two important factors are the difference between positive and negative controls and the variability within the two controls.

### Z’ was first described by J-H Zhang, TDY Chung and KR Oldenburg, 1999. A Simple Statistical Parameter for Use in Evaluation and Validation of High Throughput Screening Assays. J Biomol Screen 4:67-73.

The equation to calculate Z’ is:

**Z’ = 1 – 3*(δ_+c_ + δ_-c_)/(|μ_+c_ – μ_-c_|)**

δ_+c_ = standard deviation of the positive control

δ_-c_ = standard deviation of the negative control

μ_+c_ = mean of the positive control

μ_-c_ = mean of the negative control

Since μ_+c_ can be greater than or less than μ_-c_, depending upon the assay, the equation uses the absolute value of the difference between the means of the positive and negative control.

**Interpretations of Z’**

| **Z’** | **Interpretation** |
| --- | --- |
| 1.0 | Ideal assay. Very unlikely to achieve this |
| Between 0.5 and 1.0 | An excellent assay |
| Between 0 and 0.5 | A marginal assay |
| Less than 0 | This assay is not useful |

**SESSION 12 – developing a hypothesis and beginning experimental design**

*Recap – what were last sessions’s learning issues? What did we find out?*

***Discussion Topics***

- Any learning issues from last week that the students don’t address

*Which of your assays will you use for your high throughput screen? Your screen will produce multiple hits, i.e. drugs that test positive in your assay. What experiment will you do to decide which drug to use in your pre-clinical model?*

***Some Possible Discussion Topics***

- What would you measure if you were trying to compare two different drugs that worked by the same mechanism of action?

*Begin the design of experiments to test your compound in a pre-clinical model. What model system will you use? What measurements will you make? What controls will be necessary?*

**Some Possible Discussion Topics**

- What type of model systems are possible to use to test if your drug is effective?
- What problems can you foresee with each model system?
- What method will you use to deliver the drug?
- What could you measure to make sure the drug works *in vivo*, i.e. that it affects its target as predicted?
- What measurements could you make to determine if the drug impaired viral infection?
- What are all the controls you will need? Positive controls (does your system work)? Negative controls (is the outcome due to your drug)?

*Learning issue for Session 12 – You have modified your screen for high throughput analysis so the screen will provide a preliminary estimate of potency of each compound. The resulting data is presented in the excel spreadsheet. Which compound would you like to take into your animal model?*

*.*

**xxxxxxxxxxxxxxxxxxxxxxxxxxxxxxxxxxxxxxxxxxxxxxxxxxxxxxxxxxxxxxxxxxxxxxxxxxxxxxxxxxxx**

***Facilitator Cheat Sheet***

**Model Systems – animal models (from earlier Sessions)**

**Measurements**

**Assays for Virus**

Plaque assays on Vero E6 cells

- Serial dilutions of homogenate supernatant incubated with cells for 1 hr
- Overlay with agarose/media incubate 37oC for 2 day
- Remove overlay, stain with crystal violet and count plaques

Focus forming assay

- Serial dilutions of homogenate supernatant incubated with cells for 1 hr
- Overlay with carboxymethylcellulose incubate 24 hr
- Remove overlay, fix with paraformaldehyde and permeabilize
- Stain with antibody vs viral protein (nucleocapsid or spike)
- Count foci

qRT-PCR assay

- Could be a qPCR assay with appropriate controls and method to calculate quantity of RNA
- Could be a qPCR assay with control that can be used to generate standard curve to quantify RNA

Immunohistochemistry on tissue using viral antibodies

- Stain tissue to look for evidence of virus in different tissues

**Assays based on animal’s response to virus**

- Weight loss
- Labored breathing
- Lethargy
- Death
- Elevated inflammatory cytokines

**SESSION 13 – Therapeutics Pre-clinical Testing**

*Recap – what were last week’s learning issues? What did we find out?*

***Discussion Topics***

- Any learning issues from last week that the students don’t address

*Which drug are you going to use in your pre-clinical model?*

***Discussion Topics***

- How can you identify the best candidate from the list of drugs (lowest IC50)

*Review your model system and experimental design, including methods, measurements and controls. Are the methods and experimental approach rigorous? How could they be made more rigorous?*

***Discussion Topics***

- How many replicates or how many animals do you need for the study?
- What other methods and measures would support your experiment?

*Discuss how your controls validate the outcome of the experiment. What additional controls could increase confidence in your result?*

***Discussion Topics***

- What other positive controls could be used?
- What other negative controls could be used?

*How are you going to analyze your experimental results?*

***Discussion Topics***

- How are you going to look at your data and present your data?
- What analyses can you do to determine if the results of the controls and the experimental are truly different?

*What are your expected results and how will you interpret these results?*

***Discussion Topics***

- What do you expect to see in your measurement if the hypothesis is correct? Incorrect?
- What other explanations can result in getting the measurement you expect, yet your hypothesis is incorrect? What control might rule this out.

*If your hypothesis is supported by the results of your experiments, what will you do next?*

*If your hypothesis is not supported by your results, what modifications to your hypothesis or alternative hypothesis will you propose? How would you test this (superficial description)?*

**APPENDIX V ASSIGNMENTS**

**WRITTEN ASSIGNMENT**

Each student is responsible for working on their own to develop and complete their own writing assignment. All of the material researched by the entire team should be available to each member of the team to use to write their own assignment.

**Topic:** The assignment will be built upon the blood clotting problem, covered in Sessions 4, 5 and 6.

**Format:** Single spaced document, between 1200 and 1400 words. The document shall contain the following sections:

- Background – sufficient background to establish the basis of the hypothesis and the rationale for the proposed experiments
- Hypothesis – succinctly stated hypothesis
- Experimental plan – well designed experimental plan using multiple approaches to test the hypothesis. The plan should demonstrate rigor and include all of the essential controls to demonstrate the validity of the experiments.
- Proposed analysis – discussion of the proposed analysis of the experimental results, expected results and interpretation of your expected results.

The Written Report Rubric will be used to grade the written reports.

Your facilitator and the course coordinator will edit your assignment and provide feedback to you to help you improve your writing. It will be beneficial to you to modify your document on your own to incorporate your facilitator’s recommendations.

Unsatisfactory written assignments will receive editorial feedback from the facilitator. The assignment must be modified accordingly and an improved assignment returned to the facilitator.

**ORAL PRESENTATION ASSIGNMENT**

**Topic:** The presentation will be built upon either the neuro problem, covered in Sessions 7, 8 and 9, or the therapeutics problem, covered in Sessions 10, 11 and 12. Each team can choose which problem to develop for the oral presentation.

**Component of the presentation:** The presentation should include:

- Background – sufficient background to establish the basis of the hypothesis and the rationale for the proposed experiments
- Hypothesis – succinctly stated hypothesis and discussing the scientific premise upon which the hypothesis is based
- Experimental plan – well designed experimental plan using multiple approaches to test the hypothesis. The plan should demonstrate rigor and include all of the essential controls to demonstrate the validity of the experiments. The experimental plan should include a description of the materials and methods.
- Proposed analysis – discussion of the proposed analysis of the experimental results, expected results and interpretation of your expected results. Include a discussion of how you would proceed if your hypothesis is incorrect. Provide an alternative hypothesis and a brief outline of how you would test the hypothesis.

**Evaluation:** All team members will work together to contribute to the presentation and all team members should understand all of the parts of the presentation.

Individuals will also use the opportunity to develop their oral presentation skills. Important considerations for individual presentation skills are listed in the rubric.

**APPENDIX VI RUBRICS**

**Class participation rubric**

| **Category** | **Exemplary Performance** | **Very Good Performance** | **Satisfactory Performance** | **Needs Improvement** |
| --- | --- | --- | --- | --- |
| **Adherence to group ground rules** | Always follows the ground rules  Supports the ground rules and encourages others to do so | Always follows the ground rules with very rare lapses | Generally follows the ground rules with occasional lapses | Fails to follow the ground rules repeatedly  Flaunts the ground rules |
| **Researching learning issues** | Well researched, accurate and detailed  Demonstrates knowledge of topic | Well researched, accurate and detailed, but with some deficiencies | Deficiencies in accuracy and details | Incompletely researched, inaccuracies, lack of detail, lack of depth of knowledge |
| **Presenting issues and the problem** | Clear and accurate presentation of material, appropriate sources used, efficient summary of critical information | Clear and accurate presentation of material, appropriate sources used, missing some critical information | Some deficits in accuracy, uses questionable resources, missing critical information | Unclear presentation, inaccurate information, failure to utilize proper resources, does not cover critical information |
| **Contribution of ideas** | Consistently makes valuable contributions that stimulate discussion | Often makes valuable contributions to the discussion | Sometimes has difficulty making rational contributions to the discussion | Frequently does not make rational contributions to the discussion |
| **Creative thinking** | Comments are insightful, thinking is creative | Some comments are insightful, some thinking is creative | Difficulty in developing insightful or creative ideas | Demonstrates lack of insight or creativity in discussion of ideas |
| **Critical analysis** | Performs critical analysis, makes sound deductions, appropriate appraisal of knowledge | Often performs critical analysis, makes sound deductions, appropriate appraisal of knowledge | Deficits in critical analysis, deductions and appraisal of knowledge | Limited effort at critical thinking, does not engage in critical discussions or share ideas |
| **Professionalism Active learning** | Recognizes limitations of their knowledge, eagerly seeks additional learning | Often recognizes limitations of their knowledge, eagerly seeks additional learning | Sometimes fails to recognize the limitations of their own knowledge, hesitates to seek further knowledge | Fails to recognize the limits of their knowledge, hesitant to seek further knowledge |
| **Group interactions** | Fosters open communication, actively listens, is a positive participant in group discussions | Often fosters open communications, listens and is a positive participant in group discussions | Sometimes fails to foster open communication, is inattentive or is a negative influence in group discussions | Appears occupied when others present, monopolizes time, is a negative force within the group |
| **Participation** | Strong active participant in all activities, does their fair share, helps others achieve critical goals | Active participate in all activities, does their fair share and helps others | Sometimes refrains from participating in group activities or helping others | Limited participations, fails to do their fair share, fails to help others |
| **Group dynamics** | Seeks to focus group on task, encourages efficiency, works toward completing the task | Often seeks to focus group on task, work efficiently and toward completing the task | Often is quiet, engages in distraction that impedes progress | Distracting influence, comments are off topic, distracts from group progress |
| **Punctuality and reliability** | Always on-time and ready to contribute | Always on-time and usually ready to contribute | Always on-time but not ready to contribute | Chronically late and unprepared to participate in class |

**Written Report Rubric**

**Format:** Single spaced document, between 1200 and 1400 words. The document shall contain the following sections:

- Background – sufficient background to establish the basis of the hypothesis and the rationale for the proposed experiments
- Hypothesis – succinctly stated hypothesis
- Experimental plan – well designed experimental plan using multiple approaches to test the hypothesis. The plan should demonstrate rigor and include all of the essential controls to demonstrate the validity of the experiments.
- Proposed analysis – discussion of the proposed analysis of the experimental results, expected results and interpretation of your expected results.

| **Category** | **Exemplary Performance** | **Very Good Performance** | **Satisfactory Performance** | **Needs Improvement** |
| --- | --- | --- | --- | --- |
| **Background** | Complete, all relevant information is present, no extraneous information is present | Most or all relevant information is present, some extraneous information maybe present | Incomplete, missing relevant information, inclusion of extraneous material | Incomplete, missing many key pieces of relevant information, inclusion of large amounts of extraneous information |
| **Hypothesis** | Coherent and succinct hypothesis | Partially coherent or wordy hypothesis | Ambiguity in the hypothesis, hypothesis disconnected from background | No hypothesis |
| **Experimental plan** | Well designed experimental plan that will directly test the hypothesis, multiple approaches used, rationale for experiments built upon background | Well designed experimental plan, multiple approaches used, rationale for experiments is sound, minor deficits in some places | Some deficits in experimental design, multiple approaches used, some deficits in rationale for the experimental plan | Single experimental approach, experiment does not address the hypothesis, poor rationale for the proposed experiment |
| **Rigor** | Good incorporation of rigor into the design, limitations recognized, multiple approaches chosen to enhance rigor | Incorporates rigor, some limitations recognized, some deficits in consideration of rigor | Discusses rigor but emphasizes minor issues related to rigor, ignoring major issues in experimental design | Rigor is not considered, or is unable to distinguish pieces of experimental design requiring additional consideration of rigor |
| **Controls** | All controls present, demonstrates rationale for the controls | All controls present, missing some rationale for controls | Missing controls, extraneous controls, missing rational for controls | Major deficits in controls and rationale for controls |
| **Analysis** | Includes discussion of methods of analysis and how analysis will be performed | Includes most of the important points about methods of analysis and performance of analysis | Major deficits in discussion of analysis or how analysis will be performed | Superficial description of analysis |
| **Expected results** | Clear, rational description of all the expected results, including the results for controls and experimental | Good description of expected results but not crystal clear or partially missing a description | Only partial description of possible expected results, missing key expected results | Fail to adequately describe expected results |
| **Interpretation** | Clear, rational description of how expected results will be interpreted, includes discussion of scenario where expected results are obtained and where expected results are not obtained | Good, but incomplete description of how results will be interpreted based upon expected results | Poor description of interpretation, partial consideration of expected results, failure to connect interpretation to expected results | Superficial interpretation, incoherent interpretation |
| **Grammar** | Well written, easy to read and follow logic of the document, no grammatical errors | Well written, some points are not clear due to choice of words, few grammatical errors | Pieces are well written and other parts are difficult to understand, a number of grammatical errors | Difficult to read and comprehend, incomplete sentences, grammatical errors, poor writing obscures the meaning of the document |
| **Spelling** | No spelling mistakes | No spelling mistakes | Some typos | Many typos |

**Oral Presentation Rubric**

This will be a group presentation, ~40 minutes in length, followed by questions from the facilitator. The presentation will be built upon a problem in the course. All members of the team will participate in the presentation, which should include:

- Background – sufficient background to establish the basis of the hypothesis and the rationale for the proposed experiments
- Hypothesis – succinctly stated hypothesis and discussing the scientific premise upon which the hypothesis is based
- Experimental plan – well designed experimental plan using multiple approaches to test the hypothesis. The plan should demonstrate rigor and include all of the essential controls to demonstrate the validity of the experiments. The experimental plan should include a description of the materials and methods.
- Proposed analysis – discussion of the proposed analysis of the experimental results, expected results and interpretation of your expected results. Include a discussion of how you would proceed if your hypothesis is incorrect. Provide an alternative hypothesis and a brief outline of how you would test the hypothesis.

All team members will work together to contribute to the presentation and all team members should understand all of the parts of the presentation.

Individuals will also use the opportunity to develop their oral presentation skills. Important considerations for individual presentation skills are listed in the rubric below:

| **Category** | **Exemplary Performance** | **Good Performance** | **Needs Improvement** |
| --- | --- | --- | --- |
| **Background** | Broad and deep background providing information on all aspects of the questions. Discusses relevance/significance. Very well organized and easy to understand. Connections are very clear. | Hits the major points but misses some details. Decent organization with obvious effort. Makes the obvious connections but misses some subtle ones | Narrow and shallow background introducing only the most obvious aspects. Poor organization, difficult to follow |
| **Hypothesis** | Coherent and succinct hypothesis, connected to scientific premise | Partially coherent or wordy hypothesis | Ambiguity in the hypothesis, hypothesis disconnected from background |
| **Experimental plan** | Well designed experimental plan that will directly test the hypothesis, multiple approaches used, rationale for experiments built upon background, complete description of key reagents and experimental procedures | Well designed experimental plan, multiple approaches used, rationale for experiments is sound, deficits and lack of detail in some places | Single experimental approach, experiment does not address the hypothesis, poor rationale for the proposed experiment, lack of detail in experimental design |
| **Rigor** | Good incorporation of rigor into the design, limitations recognized, multiple approaches | Incorporates rigor, some limitations recognized, some deficits in consideration of rigor | Rigor is not considered, or is unable to distinguish pieces of experimental design requiring addition consideration of rigor |
| **Controls** | All controls present, demonstrates rationale for the controls | Most controls present, missing some rationale for controls | Major deficits in controls and rationale for controls |
| **Analysis** | Includes discussion of methods of analysis and how analysis will be performed | Some deficits in discussion of analysis or how analysis will be performed | Superficial description of analysis |
| **Expected results** | Clear, rational description of all the expected results, including the results for controls and experimental | Good description of expected results, but only partial description of possible expected results, maybe missing a key expected result | Fail to adequately describe expected results |
| **Interpretation** | Clear, rational description of how expected results will be interpreted, includes discussion of scenario where expected results are obtained and where expected results are not obtained | Good, but incomplete description of how results will be interpreted based upon expected results | Superficial interpretation, incoherent interpretation |
| **Time Management** | Presentation completed in proper time. Balanced time for different components/individuals | One or two not in sync with the rest, talk falls 5 min too short or long | Way too long, way too short. Frequently wanders far off topic |
| **Individual Presentations Skills** | | | |
| **Pace of presentation** | Evenly paced presentation that is easy to follow | Too slow at start or too fast at end, OK otherwise | Very slow or very fast |
| **Eye contact** | Constant eye contact all across the room | Engaged with audience for part but not all of presentation | Never looks at the audience |
| **Clarity** | Very clear description of all points. Very logical flow | Some areas very clear; others not well elaborated | Points glossed over or presented in a manner that is difficult to understand |
| **Aesthetics (if applicable)** | Highly organized, very easy to read/understand, crystal clear | Some slides look excellent while others are hard to decipher | Material is difficult to read and disorganized |
| **Grammar** | No major errors | Some errors but mostly minor | Major grammatical errors |
| **Ability to answer questions (if applicable)** | Student is knowledgeable, answers questions intelligently | Student answers some questions well and thoroughly, misses the mark on others | Student does not appear to have a clear understanding |
| **Accuracy** | Few minor errors in fact, if any | Some minor fact errors, but major points accurate | Major errors in facts |

**Peer to Peer Evaluation Rubric**

The grading scale and rubric are based on a Goldilocks scale. The “best” grade is in the middle and poorer scores are on each end.

| **Category** | **Too Much** | **Just Right** | **Too Little** |
| --- | --- | --- | --- |
| **Accountability** | Controlling: excessive fault finding: self-righteous | Punctual to class or duty; well prepared: willing to accept praise or constructive evaluations: follows up on communications \| | Doesn’t complete tasks on time: misses appointments: avoids work or responsibility |
| **Responsiblity** | Inflexible: rule-bound to the point of obstruction or paralysis: afraid to act out of fear of committing an error: assumes blame inappropriately: overly reliant on rules | Reliable; trustworthy: takes ownership of assignments: seriously and diligently works on assigned tasks: Wears appropriate protective clothing and gear | Inflexible: rule-bound to the point of obstruction or paralysis: afraid to act out of fear of committing an error: assumes blame inappropriately: overly reliant on rules |
| **Respectful and Nonjudgmental Behavior** | Excessive selflessness: overextends oneself; nonjudgmental to the point of inaction | Consistently civil and courteous to all: tolerates diversity: listens before acting: considers others’ feelings, background and perspectives | Little compassion for others; appears cold, heartless, indifferent |
| **Compassion and Empathy** | Loses objectivity by excessive desire to help: emotionally labile and unduly empathetic; | Respects & is aware of others’ feeling: can put oneself in other’s place: shows mindfulness & self-reflection | Little compassion for others; appears cold, heartless, indifferent |
| **Maturity** | Puts others ahead of self to a fault: attempts to improve oneself to a fault; perfectionist | \| Shows personal growth; recognizes & correct mistakes; tries to improve self; manages relationships & conflicts well; seeks feedback & modifies behavior accordingly; maintains appropriate public dress/appearance | Makes unsound decisions: doesn’t manage time well; unable to see big picture: can’t maintain personal or professional boundaries: blames others |
| **Communication** | Gives feedback when not solicited; constantly assumes role of conflict manager; self-professed communication skills are overstated | Effective use of oral, written & non-verbal skills: speaks with clarity to all; culturally appropriate skills; gives & receives constructive criticism | Unable to communicate at another’s level of understanding: not seek feedback that he/she is understood; writes illegibly; wears inappropriate dress |
| **Self-Directed Learning and Appraisal** | Dominant, overbearing, authoritative in team settings: Constantly creating change to the point of disruption | Displays ability to be a life long learner; completes all evaluations and academic and clinical work in a truthful, thoughtful & timely manner | Accomplishes tasks with excessive assistance of others: Doesn’t work well in team situations: not open to change |

**APPENDIX VII EVALUATION SURVEYS**

**Course Evaluation**

Q1. The course was well organized. Likert scale – 1 = strongly disagree, 5 = strongly agree.

Q2. Participation in this course helped me become an active learner. Likert scale – 1 = strongly disagree, 5 = strongly agree.

Q3. Participation in this course motivated me to learn. Likert scale – 1 = strongly disagree, 5 = strongly agree.

Q4. The learning issues generated in my group stimulated me to locate and interpret scientific literature. Likert scale – 1 = strongly disagree, 5 = strongly agree.

Q5. Participation in this course helped develop my problem-solving skills. Likert scale – 1 = strongly disagree, 5 = strongly agree.

Q6. Participation in this course helped me develop skills in hypothesis testing and experimental design. Likert scale – 1 = strongly disagree, 5 = strongly agree.

Q7. Participation in this course helped develop my writing skills. Likert scale – 1 = strongly disagree, 5 = strongly agree.

Q8. Participation in this course helped develop my oral presentation skills. Likert scale – 1 = strongly disagree, 5 = strongly agree.

Q9. Participation in this course helped develop my teamwork skills. Likert scale – 1 = strongly disagree, 5 = strongly agree.

Q10. The overall education value of this course is high. Likert scale – 1 = strongly disagree, 5 = strongly agree.

**Evaluation of Facilitators**

Q1. The facilitator did a very good job guiding the group by asking questions. Likert scale – 1 = strongly disagree, 5 = strongly agree.

Q2. The facilitator helped the group to stay “on track”. Likert scale – 1 = strongly disagree, 5 = strongly agree.

Q3. The facilitator willingly accepted feedback from the group and was non-defensive. Likert scale – 1 = strongly disagree, 5 = strongly agree.

Q4. The facilitator was very enthusiastic about the group. Likert scale – 1 = strongly disagree, 5 = strongly agree.

Q5. The facilitator helped the group bond as a team. Likert scale – 1 = strongly disagree, 5 = strongly agree.

Q6. The facilitator helped identify gaps in the groups’ knowledge. Likert scale – 1 = strongly disagree, 5 = strongly agree.

Q7. The facilitator helped the group set appropriate learning issues. Likert scale – 1 = strongly disagree, 5 = strongly agree.

**Peer to peer evaluation survey (Goldilocks scale)**

**Consult the Peer to Peer Evaluation Rubric to appropriately evaluate**

This evaluation is intended to provide insight into professionalism and teamwork in BMS706. The scale is bi-polar, meaning that a student may exhibit too much (closer to 7) or too little (closer to 1) of the preferred professional/team behavior. A “4” is the preferred behavior, which characterizes a student who exhibits the appropriate professional behavior.

Q1. Accountability

0 = not observed

1 = not accountable, e.g. always late

4 = accountable, e.g. punctual, well prepared, follows up

7 = controlling, e.g. finds fault in others

Q2. Responsibility

0 = not observed

1 = irresponsible, e.g. breaks rules, makes excuses, deflects blame

4 = responsible, e.g. reliable, diligent, takes ownership

7 = inflexible, e.g. overly reliant on rules, afraid to act, assumes blame inappropriately

Q3. Respectful and nonjudgmental behavior

0 = not observed

1 = disrespectful and judgmental, e.g. makes assumptions, discourteous, belittling

4 = respectful and nonjudgmental, e.g. civil, courteous, tolerant, listens, empathetic

7 = excessively selfish or nonjudgmental to the point of inaction

Q4. Compassion and empathy

0 = not observed

1 = incompassionate, e.g. appears cold, heartless or indifferent

4 = compassionate, e.g. respects others, empathetic, mindful, self-reflective

7 = excessive desire to help, emotionally labile, unduly empathetic

Q5. Maturity

0 = not observed

1 = immature, e.g. unsound decisions, poor time management, blames others

4 = mature, e.g. recognizes and corrects own mistakes, tries to improve, manages relationships

7 = perfectionist, e.g. puts others ahead of self to a fault, tries to improve to a fault

Q6. Communication

0 = not observed

1 = poor communication, e.g. can’t communicate to others

4 = good communication, e.g. effective oral communication, clear, gives and receives constructive criticism

7 = gives unsolicited feedback, always assumes role of conflict manager, self-professed skills are overrated

Q7. Self-directed learning and appraisal

0 = not observed

1 = overly dependent, e.g. completes tasks with excessive assistance, poor teamwork

4 = balance, e.g. displays self-directed learning and works well with the team to share knowledge

7 = controlling, e.g. authoritative in team settings, constantly trying to change things to the point of disruption

Q8. The student followed the rules established by the group. Likert scale – 1 = strongly disagree, 5 = strongly agree.

Q9. I would want to have this student as a member of my team. Likert scale – 1 = strongly disagree, 5 = strongly agree.

**APPENDIX VIII GRADING RUBRIC**

Grading key (0 points – 16 points)

Choosing a model system to address the scientific problem: cell line or model organism

a. Model system irrelevant for the scientific question (0 points)
b. Model system will work but is not described in sufficient detail (1 point)
c. The model system proposed is adequate for the experiment and accurately described, demonstrating good rationale to test the hypothesis (2 points)
d. Alternative model system is proposed (3 points)

Description of the readout of the experiment proposed

a. Readout is not appropriate (0 points)
b. Readout is appropriate but not sufficiently described (1 point)
c. Correct readout, accurately described (2 points)
d. Alternative experiment with a different readout proposed (3 points)

Description of the experimental controls

a. No controls suggested (0 points)
b. Controls suggested but inadequate for the experiment proposed (Either positive or negative control) (1 point)
c. Both positive and negative control described (2 points)
d. Additional controls suggested (3 points)

Describe the expected results and how you would analyze the data

a. Description of expected results and analysis is not included (0 points)
b. Description of expected results and analysis is included but doesn’t match the experiment or the scientific problem (1 point)
c. Description of expected results and analysis is included with sufficient details, demonstrating good rationale for the approach (2 points)
d. Additional/alternative experiments with expected results are described, including alternative if the results don’t support the hypothesis (3 points)

Statistical significance

1. There is no consideration about statistical significance (0 points)
2. Attempt to address statistical significance but insufficient details/wrong numbers (1 point)
3. Correct and adequate description of how statistical significance will be achieved – demonstrating rationale for analysis of the data (2 points)!

Clarity of the text

1. Difficult to read or key details missing (0 points)
2. Description shows general understanding of the scientific problem but lacks scientific terms and some details (1 point)
3. Description shows depth in knowledge, follows scientific logic; scientific terms properly used (2 points)
